# Mouse genomic rewriting and tailoring: synthetic *Trp53* and humanized *ACE2*

**DOI:** 10.1101/2022.06.22.495814

**Authors:** Weimin Zhang, Ilona Golynker, Ran Brosh, Aleksandra M. Wudzinska, Yinan Zhu, Lucia Carrau, Payal Damani-Yokota, Camille Khairallah, Noor Chalhoub, Emily Huang, Hannah Ashe, Kamal M. Khanna, Matthew T. Maurano, Sang Yong Kim, Benjamin R. tenOever, Jef D. Boeke

## Abstract

Genetically Engineered Mouse Models (GEMMs) aid in understanding human pathologies and developing new therapeutics, yet recapitulating human diseases authentically in mice is challenging to design and execute. Advances in genomics have highlighted the importance of non-coding regulatory genome sequences controlling spatiotemporal gene expression patterns and splicing to human diseases. It is thus apparent that including regulatory genomic regions during the engineering of GEMMs is highly preferable for disease modeling, with the prerequisite of large-scale genome engineering ability. Existing genome engineering methods have limits on the size and efficiency of DNA delivery, hampering routine creation of highly informative GEMMs. Here, we describe mSwAP-In (<u>m</u>ammalian <u>Sw</u>itching <u>A</u>ntibiotic resistance markers <u>P</u>rogressively for <u>In</u>tegration), a method for efficient genome rewriting in mouse embryonic stem cells. We first demonstrated the use of mSwAP-In for iterative genome rewriting of up to 115 kb of the *Trp53* locus, as well as for genomic humanization of up to 180 kb *ACE2* locus in response to the COVID-19 pandemic. Second, we showed the *hACE2* GEMM authentically recapitulated human *ACE2* expression patterns and splicing, and importantly, presented milder symptoms without mortality when challenged with SARS-CoV-2 compared to the *K18-ACE2* model, thus representing a more authentic model of infection.

## Introduction

Genome synthesis is feasible for some prokaryotes such as *Escherichia coli*^1^, *Mycoplasma*^2, 3^, and eukaryotes such as *Saccharomyces cerevisiae*^4–11^. However, mammalian genome synthesis is still prohibitive due to the enormous genome size and complexity^12^. A bold step towards mammalian genome writing is to overwrite large swaths of a native genomic region that covers a full gene, complete with all regulatory regions, or even several nearby genes. The combination of big DNA assembly approaches with site-specific recombinases in mammalian systems has proven to be an efficient way to modify mammalian genomes in a large-scale fashion^13–16^. However, limitations of most existing big DNA delivery methods must still be overcome as present technologies leave significant scars behind in the genome^15^, although this problem is largely solved by the recently developed Big-IN method^14^, but methods are usually not designed for iterative deliveries, limiting delivery size of incoming DNA. A cleaner, more efficient mammalian genome writing method that can in theory be used to overwrite entire mammalian chromosomes will benefit a wide variety of functional studies.

Mouse embryonic stem cells (mESCs) are relatively easy to genetically manipulate and subsequent derivation of mouse models is possible by the generation of chimeras or by tetraploid complementation^17–20^. Although efforts have been made to create many GEMMs to model different diseases, routinely creating highly informative GEMMs has still failed to reach its potential due to the lack of reliable genomic tools. Genetically humanizing mouse loci bridges human-mouse evolutionary gaps, reflected in some cases by the lack of clear-cut human orthologs^21^ and the failure to recapitulate human disease^22–24^. In the past several decades, transgenesis has been the predominant approach for mouse humanization, a process which typically delivers the coding sequence of human genes under a strong exogenous promoter to gain the biochemical characteristics of human proteins, but at the expense of non-physiological expression patterns. Projects like ENCODE^25^ and GWAS^26^ highlight the importance of regulatory elements and single nucleotide polymorphisms in non-coding regions, making full genomic humanization (including non-coding regions) preferable. Human YAC/BAC-based transgenes can retain the full-length sequence of human genes, but are often randomly integrated into the mouse genome^27–29^, leading to poorly-characterized position effects. This approach is therefore unable to reliably mimic the endogenous genomic context and thus compromises the expression authenticity of human genes. Precision tailoring of the YAC/BACs and the *in situ* rewriting of the mouse counterpart(s) represents excellent strategies for overcoming the drawbacks of YAC/BAC-based transgenes. Previous work on *in situ* humanization of 6 Mb of mouse immunoglobulin genes set a good example. However, the overall efficiency for each human sequence integration in those methods was no higher than 0.5%^30, 31^, limiting the widespread application of the method.

During the COVID-19 pandemic, one of the many substantial challenges faced was the inability to use the mouse as a small animal model to understand the disease. Due to natural polymorphisms in murine Ace2, the receptor for SARS-CoV-2, the original isolates of the virus such as the Washington strain, are unable to productively infect mice. While the virus can be adapted to mice^32, 33^, studying the biology of a modified virus limits the value of what can be learned from such a model. Similarly, current animal models in which human *ACE2* is genetically introduced to mice, e.g. by driving expression with a strong promoter^24^ often leads to changes in viral tropism not observed physiologically. While recent variants of SARS-CoV-2 have gained the capacity to infect mice^34^, the host response fails to phenocopy what is observed in humans^35^. Given this, a mouse model that is susceptible to SARS-CoV-2 and has the ability to mimic human disease pathology could be extremely valuable for therapeutic development, as well as for a better basic understanding of human COVID pathophysiology, and the effects of age, immune-suppression status and other factors. Such a model could leverage the enormous genetic resources available for murine-based models. Such models are also valuable for preparedness against potential future disease outbreaks. The transgenic *ACE2* mouse models that were developed in response to previous SARS-CoV and MERS-CoV outbreaks provided great platforms for understanding these diseases^24, 36–38^. Yet, the existing transgenic *ACE2* mouse models have several limitations: First, because they lack the human regulatory elements around *ACE2*, they are consequently unable to recapitulate the spatiotemporal regulation of human *ACE2*. For example, human *ACE2* is strongly expressed in the testis, whereas mouse *Ace2* does not. Second, the mouse gene likely lacks the alternative splicing elements required to produce certain human-specific isoforms^39^. Third, leaving the mouse endogenous *Ace2* gene intact results in a complicated and uncertain mixture of both human and mouse ACE2 proteins. A genomically humanized *ACE2* mouse that more accurately models coronavirus diseases is sorely needed.

Here we first report a novel mammalian genome writing method, mSwAP-In, for precise, efficient, scarless, and iterative genome writing in mESCs. Prior to the pandemic, we developed mSwAP-In to address an earlier gauntlet thrown down by the Genome Project-Write project^40^: to engineer a synthetic *Trp53* with recoded mutational hotspots, which might render cells more resistant to spontaneous oncogenic *Trp53* mutations. We used this platform to highlight mSwAP-In’s utility for delivery of a synthetic mouse gene, as well as for iterative genome writing by efficiently overwriting the regions downstream of *Trp53* with three carefully designed secondary payload DNAs. To generate a humanized mouse model recapitulating COVID-19 pathology, we efficiently swapped 72 kb of the mouse *Ace2* locus with 116 kb or 180 kb of the human *ACE2* genomic region. The subsequently-generated *hACE2* GREAT-GEMM (Genomically Rewritten and Tailored GEMM) accurately reflected human-specific aspects of authentic gene expression both at the transcriptional and splicing levels. *ACE2*-humanized mice were susceptible to SARS-CoV-2 upon intranasal infection, but unlike the transgenic *K18-hACE2* model, the animals did not succumb to infection, suggesting that the *hACE2* GREAT-GEMM may better model human COVID-19.

## Results

### Design of mSwAP-In

Most genome engineering methods are restricted by difficulties in DNA assembly, purification and delivery to mammalian cells as construct length increases. To overcome the size limitation, we designed <u>m</u>ammalian <u>Sw</u>itching <u>A</u>ntibiotic markers <u>P</u>rogressively for <u>In</u>tegration (mSwAP-In), a method directly descended from our proven-effective yeast genome rewriting method, SwAP-In^4, 6^. Like SwAP-In, mSwAP-In is designed to overwrite hundreds of kilobases of wild-type mammalian genome segments with synthetic DNA in a scarless and iterative manner. In theory, mSwAP-In, like SwAP-In, could be used to overwrite an entire chromosome by iteration, although hundreds of such steps would be required to do this for even the smallest mammalian chromosome.

Two marker cassettes (MC1 and MC2) were designed to deploy mSwAP-In (**Fig. 1a**). Each consists of a distinct set comprising: 1) a fluorescence marker serving as an indicator of positive clones during or after colony picking; 2) a positive selection marker; and 3) a negative selection marker that is overwritten together with wild-type DNA in each swapping step, allowing selecting against off-target integrations. A series of marker cassettes are designed to accommodate genetic backgrounds that already contain selectable markers (**Fig. S1a**). Each selection component was tested for effective elimination of sensitive mESCs (**Fig. S1b**). Finally, a universal gRNA target (UGT) site orthogonal to mammalian genomes (derived from GFP) was placed in front of each marker cassette to allow specific and efficient cleavage by Cas9-gRNAs or other nucleases. To enable use of the *HPRT1* minigene in MC2 in later mSwAP-In stages, mESCs were pre-engineered to delete the endogenous *Hprt* gene using two Cas9-gRNAs followed by 6-Thioguanine (6-TG) selection^41^ (**Fig. S1c**). mSwAP-In is executed in several steps: 1) MC1 is inserted at a “safe” location near the genomic region of interest using CRISPR-Cas9 assisted homologous recombination (**Fig. 1b, Step 1**). 2) A synthetic payload DNA consisting of flanking UGT1 sites, homology arms (HAs, ∼2 kb at each end) and MC2 is pre-assembled in yeast^42^, and then co-delivered with two Cas9-gRNAs recognizing UGT1 and the distal boundary of the native genomic segment to be overwritten (**Fig. 1b, Step 2**). Payload DNA integration by homologous recombination (HR) is assisted by linearization of payload DNA at two flanking UGT1 sites and by DNA double strand breaks at targeted genomic region. Cells in which targeting was successful are selected for the presence of MC2’s positive selection marker (BSD) and parental MC1’s negative selection marker (ΔTK) (**Fig. S1d**), resulting in the wild-type segment being overwritten by the synthetic payload DNA. Iterating this process in Step 3 with a second synthetic payload DNA, assembled similarly in yeast with HAs and MC1, is performed by positively selecting MC1’s PuroR and negatively selecting against MC2’s *HPRT1* (**Fig. 1b, Step 3**). The iteration can in principle continue indefinitely as needed. Once the writing is finished, the last marker cassette can be removed either by employing CRISPR-Cas9 assisted HR or by a PiggyBAC excision system^43^, and scarlessly-engineered cells can be isolated using negative selection (**Fig. 1b, Step 4; Fig. S1a**).

**Fig. 1.**
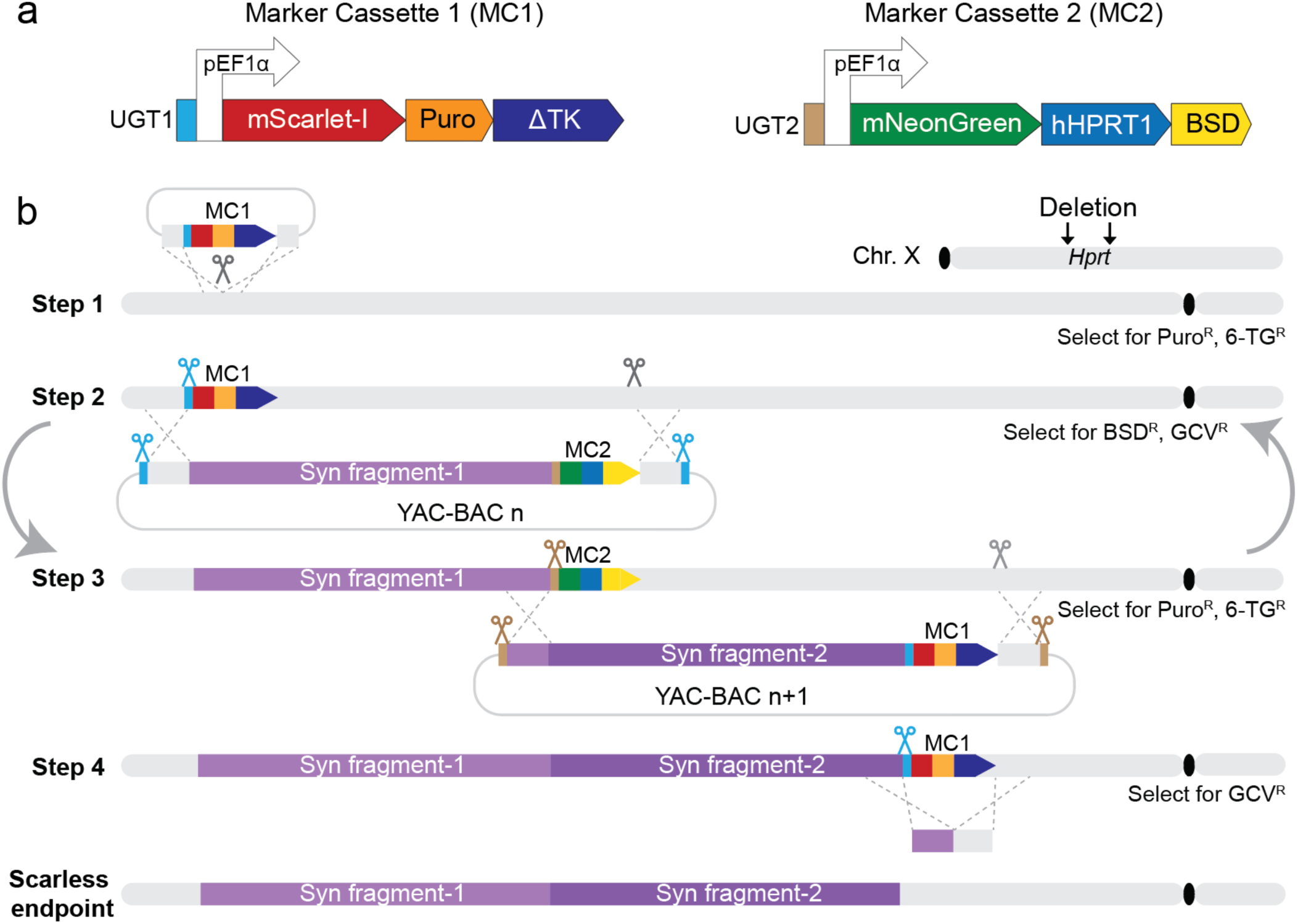
mSwAP-In strategy for genome writing. **(a)** Two interchangeable marker cassettes (MC) underlie mSwAP-In selection and counterselection. UGT, universal gRNA target; Puro, puromycin-resistance gene; BSD, blasticidin S deaminase; ΔTK, a truncated version of herpes simplex virus type 1 thymidine kinase. **(b)** Stepwise genome rewriting using mSwAP-In. A prior engineering step to delete endogenous *Hprt* enables later iteration. Step 1: integrate marker cassette 1 upstream of locus of interest. Step 2: deliver payload DNA and Cas9-gRNAs for integration through homologous recombination. Step 3: deliver next payload DNA following same strategy as Step 2, swapping back to marker cassette 1. Iterative Steps 2 and 3 can be repeated indefinitely using a series of synthetic payloads, by alternating selection for marker cassettes (curved arrows). Step 4: remove final marker cassette. YAC: yeast artificial chromosome, BAC: bacterial artificial chromosome, 6-TG: 6-Thioguanine, GCV: ganciclovir. Gray bars are native chromosome regions, purple bars are synthetic incoming DNAs. Blue and brown scissors are universal Cas9-gRNAs cutting UGT1 and UGT2, respectively; gray scissors are genomic-targeting Cas9-gRNAs. Superscript R, drug resistance.

### Rewriting *Trp53* locus with mSwAP-In in mESCs

We initially sought to engineer a “cancer-mutation-resistant” *Trp53* (p53) gene^40^ in mESCs by genome writing. Missense mutations occur frequently in p53 in cancer cells and are concentrated in its DNA binding domain, at CG sites^44–46^. This is due to frequent deamination of 5-methylcytosine at CG sites leading to C to T (or G-A on the antisense strand) conversion^47, 48^, as well as the binding of DNA adducts to certain methylated CG dinucleotides^49–51^. Given that the methylated CG dinucleotides are highly mutable, we hypothesized that synonymously recoding the DNA sequence of p53 to avoid CG dinucleotides will minimize its mutation rate. To mitigate the risk of affecting the methylation landscape, only the CG dinucleotides at p53 mutation hotspots (R172, R245, R246, R270, R279) were recoded to AG (**Fig. 2a, Fig. S2a**).

**Fig. 2.**
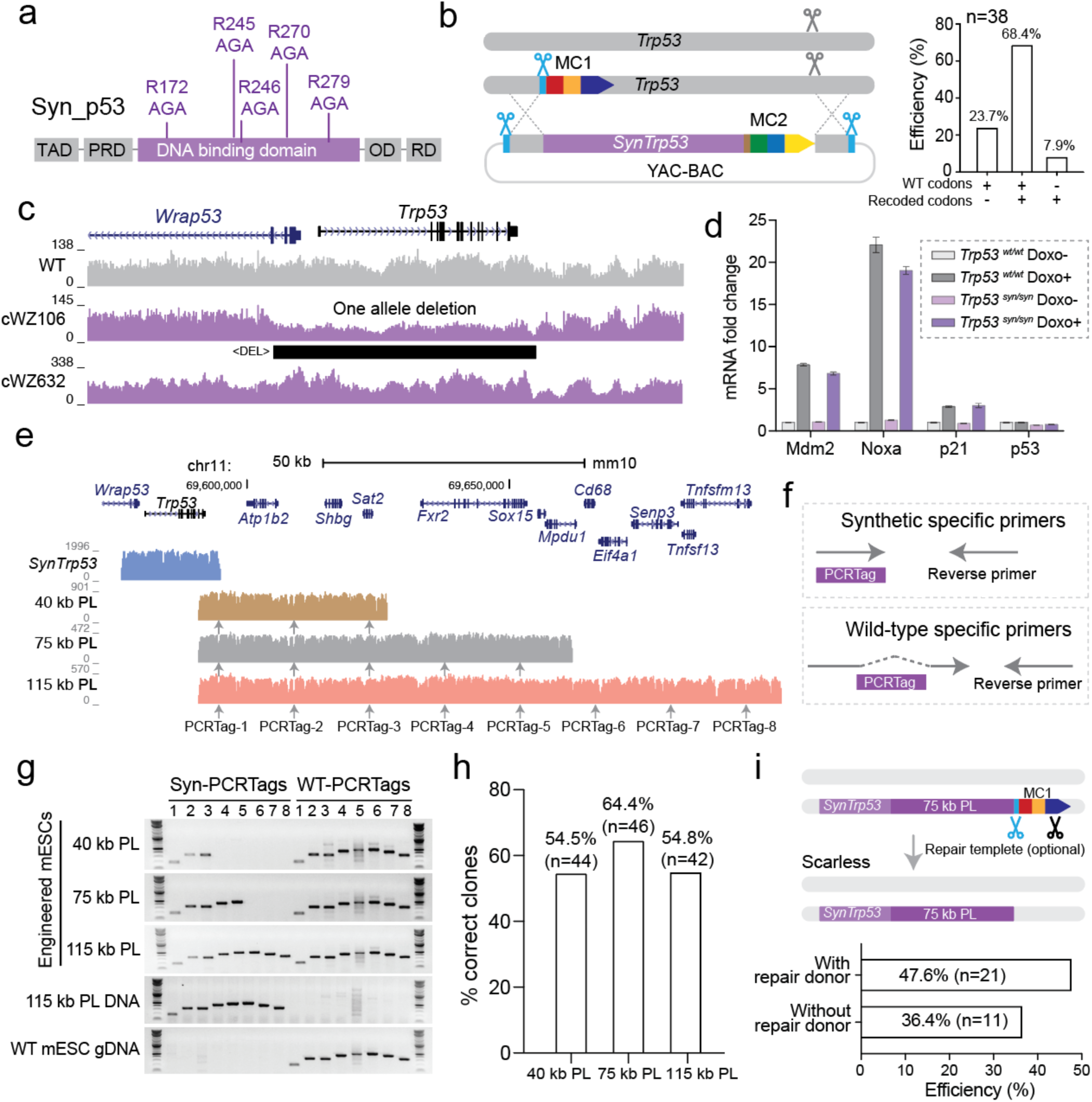
Rewriting *Trp53* locus with mSwAP-In. **(a)** Design of p53 hotspot mutation recoding. Recoded codons are shown on top. TAD, transactivation domain; PRD, proline-rich domain; OD, oligomerization domain; RD, regulatory domain. **(b)** Schematic of *synTrp53* mSwAP-In and summary of efficiency. **(c)** Sequencing coverage for wild-type, hemizygous and heterozygous clones. Sequencing reads were mapped to mm10. Black bar indicates a deletion called by DELLY^53^ **(d)** Functional evaluation of recoded p53. mESCs with wild-type or CG-recoded *Trp53* gene were treated with 250 nM of doxorubicin for 20 hours. *Mdm2*, *Pmaip1*, *Cdkn1a*, and *Trp53* mRNA levels were evaluated via RT-qPCR. mRNA levels were normalized to *mActb*, bars represent mean ± SD of three technical replicates. **(e)** Sequence coverage of *synTrp53* and three *Trp53* downstream payloads (PL) aligned to mm10. Gray arrows indicate the position of PCRTags. **(f)** Synthetic- and wild-type-specific PCR assays employing a specific forward primer and a universal reverse primer. **(g)** Genotyping of the three representative mSwAP-In integrants from three *Trp53* downstream payloads. **(h)** Summary of mSwAP-In success rates based on genotyping. **(i)** Final marker cassette removal strategy and genotyping-based efficiency summary. Blue scissors indicate UGT1-targeting gRNA; black scissors indicate gRNA targeting the SV40 terminator.

For assembly of the synthetic *Trp53* (*synTrp53*) gene, a total of 18.7 kb wild-type *Trp53* region was segmented into small DNA fragments, and the recoding sites were introduced using PCR primers. Overlapping *Trp53* fragments, linker DNAs containing the UGT1 site and MC2 were assembled in yeast (**Fig. S2b**). The successful “assemblons”^16^ were verified by restriction enzyme digestions and next-gen sequencing (**Fig. S2c-d**). In parallel, we inserted MC1 downstream of mouse *Trp53* heterozygously (**Fig. S2e**), generating an MC1-founder mESC line for *synTrp53* mSwAP-In. After deploying mSwAP-In, we found that 87.1% of the colonies lost MC1 and gained MC2 by performing genotyping PCR (n=132). We Sanger sequenced 38 genotype-verified clones and discovered that 26 of them carried the recoded codons in one of the two alleles, 9 of them were unedited, and surprisingly, 3 of them only carried recoded codons (**Fig. 2b, Fig. S2f**). *Trp53* copy number analysis of those 3 clones carrying recoded codons only suggested they were hemizygous, i.e. had only one allele (**Fig. S2g**), which was confirmed by targeted resequencing (Capture-seq^14^) (**Fig. 2c**). To ensure mSwAP-In engineering was free of off-targeting, we implemented our previously developed *bamintersect* analysis^14^, a modular mapping tool that detects reads spanning two references (e.g. payload DNA vs. mm10, homology arm vs. mm10). This analysis detected no off-target junctions in the six sequenced clones, but we detected YAC/BAC backbone integration in one clone (**Supplementary file 1**). *SynTrp53* heterozygotes can be further engineered to homozygotes by repeating mSwAP-In on the wild-type allele, but using a different version of MC2 (**Fig. S1a**).To test *synTrp53* function, we treated a homozygous *synTrp53* mESC line (*Trp53^syn/syn^*) and wild type (*Trp5^wt/wt^*) with doxorubicin, which induces DNA damage by intercalation^52^. We found that three classic p53 target genes *Mdm2*, *Pmaip1* (Noxa) and *Cdkn1a* (p21) were upregulated upon doxorubicin treatment in *Trp53^syn/syn^*mESCs to a similar degree as in wild-type mESCs, suggesting that recoding of *Trp53* did not impair its transactivation function (**Fig. 2d**).

To demonstrate the iterability of mSwAP-In and to probe the upper genome writing length limit of each mSwAP-In step, we built 40 kb, 75 kb and 115 kb payload constructs using the DNA sequence downstream of *Trp53* for the second round of mSwAP-In (**Fig. 2e, Fig. S3a**). To distinguish synthetic DNA from wild type, we inserted watermarks evenly distributed across the constructs (every ∼13 kb in intronic or intergenic regions); we refer to these watermarks as “PCRTags”, which are 28 bp orthogonal DNA sequences (**Table S1**), reminiscent of the PCRTags used in the Synthetic Yeast Genome Project (Sc2.0)^6^. Taking advantage of these PCRTags, we designed synthetic- or wild-type-specific primer pairs (**Fig. 2f**). After deploying mSwAP-In into a heterozygous *synTrp53* mESC clone, we observed the gain of synthetic PCRTags for the delivered payloads, as well as the wild-type PCRTags, indicating heterozygous integration (**Fig. 2g**). Although the total drug resistant colony number decreased as the length of the payload increased (**Fig. S3b**), the efficiency of mSwAP-In remained above 50% (**Fig. 2h**).

Lastly, we demonstrated the feasibility of marker cassette removal. Two Cas9-gRNA plasmids were used to cut at the UGT1 site and at the SV40 terminator, followed by ganciclovir counterselection. We found that MC1 removal efficiency was 47.6% when providing a ∼2 kb repair template, and 36.4% when no repair template was provided (**Fig. 2i, Fig. S3c**). Collectively, these data highlight mSwAP-In as an efficient method for large-scale iterative and scarless genome rewriting in mESCs. However, all the payloads we delivered so far are >99% identical to the native mouse genome, which might have contributed to the high efficiency. Next, we asked whether mSwAP-In could be used to overwrite the native genome with nonhomologous DNA such as entire human loci.

### Fully humanizing *ACE2* in mESCs

Mice are not naturally susceptible to SARS-CoV-2 due to differences in key residues of ACE2 required for interaction with the Spike protein^54–56^. However, the *K18-hACE2* transgenic mouse, in which a keratin 18 promoter drives high levels of expression of a human *ACE2* CDS in epithelial tissues, including respiratory epithelia, is readily infected^24^, but unlike the case of the human infection, ∼100% of the infected mice succumb to infection within days as a result of viral encephalitis^57^ – a phenotype not observed in humans. To establish a more physiological model, we aimed to completely swap the mouse *Ace2* (*mAce2*) locus with *hACE2* including all introns and regulatory elements using mSwAP-In (**Fig. 3a, Fig. S4a**). Based on the gene annotation, we noticed a long transcript (NM_001386259.1, also known as transcript variant 3) that spans 83 kb and largely overlaps with the *BMX* gene (**Fig. 3a**). In contrast to the canonical transcript which encodes an 805-aa protein, the long transcript encodes a 786-aa ACE2 protein lacking an intact collectrin homology domain at the C terminus and instead includes a novel 16-aa exon^58^. To retain all possible functions, we defined the left boundary of the payloads to include this putative long transcript. For the right boundary, considering DNase hypersensitive sites and H3K27 acetylation marks (ENCODE), we designed two *hACE2* payloads: one extending to the 3’ end of the *CLTRN* gene (*hACE2* payload 1, 116 kb), and the other one extending to the 5’ end of *CLTRN* gene (*hACE2* payload 2, 180 kb) (**Fig. 3a**).

**Fig. 3.**
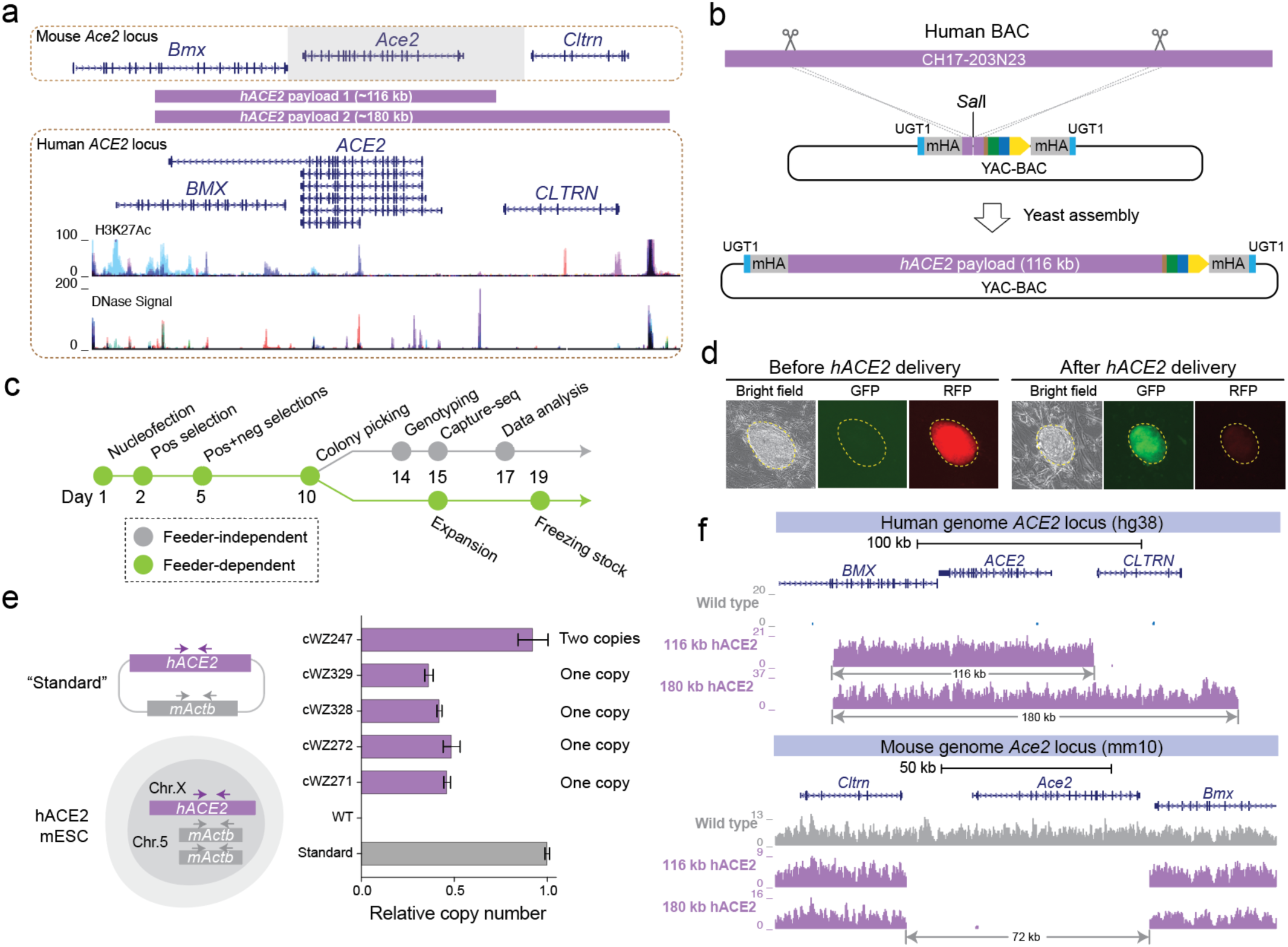
Fully humanizing *ACE2* in mESCs. **(a)** Browser shots of *mAce2* and *hACE2* loci. H3K27 acetylation and DNase signal tracks (ENCODE) in *hACE2* locus indicate functional regulatory elements. Gray box demarcates overwritten mouse genomic region. Purple bars demarcate human genomic regions included in the *hACE2* payloads. **(b)** *hACE2* payload assembly strategy. Scissors mark *in vitro* CRISPR-Cas9 digestion sites. mHA, mouse homology arm. **(c)** mESC engineering workflow. **(d)** Representative images of fluorescence marker switching in outlined mESC clones. **(e)** *hACE2* copy number determination by qPCR. The ratio between *hACE2* and *mActb* is 0.5, indicating a single copy of *ACE2* was delivered to the male mESCs, as expected. Copy number was normalized to the *mActb* gene. Bars represent mean ± SD of three technical replicates. **(f)** Sequencing coverage of 116 kb *hACE2* and 180 kb *hACE2* mSwAP-In clones. Reads were mapped to hg38 (up) and mm10 (bottom).

In contrast to the previous payload assembly strategy, the 116 kb *hACE2* region was released from a human BAC (CH17-203N23) via *in vitro* Cas9-gRNA digestion, and assembled through yeast HR into an acceptor vector^14^ that contained flanking UGT1 site, left and right *mAce2* homology arms, and MC2 (**Fig. 3b**). Correct assemblons were verified by restriction enzyme digestions followed by pulse-field gel electrophoresis (**Fig. S4b**). *hACE2* payload 2 (180 kb) was built by inserting an additional ∼64 kb fragment released from another human BAC (CH17-449P15) into the end of *hACE2* payload 1 (**Fig. S4c**). Sequencing revealed no variants within the two payloads except single nucleotide polymorphisms (SNPs) that originated from the parental BACs, highlighting the high accuracy of this BAC-based big DNA assembly workflow (**Fig. S4d**). To enable *hACE2* payloads delivery we inserted MC1 downstream of *mAce2* in C57BL/6J mESCs (**Fig. S4e**). We used feeder-dependent cell culture conditions to maintain the developmental potential of the mESCs, while splitting cells from each clone into a feeder-independent subculture for genotyping and sequencing (**Fig. 3c**). We delivered both *hACE2* payloads into the MC1 founder line, applied positive and negative selections sequentially, and observed the switch of fluorescent markers from red to green (**Fig. 3d**), indicating a successful marker cassette swap. To ensure the *mAce2* locus was fully overwritten by the two *hACE2* payloads, we performed genotyping PCR using multiple primers across *mAce2* and *hACE2* regions. Correct clones showed the presence of *hACE2* amplicons and the absence of *mAce2* amplicons (**Fig. S4f**). The overall efficiency was 61.5% (n=13) for the 116 kb *hACE2* payload, and 60.8% (n=79) for the 180 kb *hACE2* payload as determined by genotyping PCR, which is >50 times higher than previously reported methods^30, 59^.

To enable *hACE2* copy number quantification, we constructed a plasmid containing one copy of *mActb* and one copy of *hACE2* to serve as a standard in qPCR analysis, and identified mESC clones with one copy of *hACE2* (**Fig. 3e**). Lastly, Capture-seq analysis verified that the *hACE2* mESC clones have even coverage across the *ACE2* region with no deletion or duplication events, as well as the loss of mouse *Ace2* (**Fig. 3f**). We also confirmed a lack of off-target integration for *hACE2* by *bamintersect* analysis (**Supplementary file 1**), and no Cas9 reads were captured in these mESC clones (**Fig. S4g**). Considering all the steps of this comprehensive sequence quality control, the overall success rates for the 116 kb and 180 kb *hACE2* payloads were 15.4% and 22.8%, respectively.

### *hACE2* mice display physiological *ACE2* expression and splicing

*hACE2* mESCs that passed our stringent verification pipeline were subjected to blastocyst embryo injection and tetraploid-blastocyst embryo injection, which requires more naïve developmental pluripotency^17–19^. Coat color chimerism was observed with high efficiency (31 of 45 pups) when injecting the 116 kb *hACE2* mESCs into wild-type blastocysts (**Fig. 4a**). Indeed, some of the chimeric males showed 100% germline transmission (**Fig. S5a**). When injecting the 116 kb *hACE2* and 180 kb *hACE2* mESCs into a tetraploid blastocyst for embryo complementation, 14% (n=50) and 22.9% (n=70) birth rates were observed, respectively (**Table S2**). We genotyped various tissues from a tetraploid complementation-derived mouse, and detected only *hACE2* amplicons, indicating the mice were purely developed from *hACE2* mESCs (**Fig. S5b**).

**Fig. 4.**
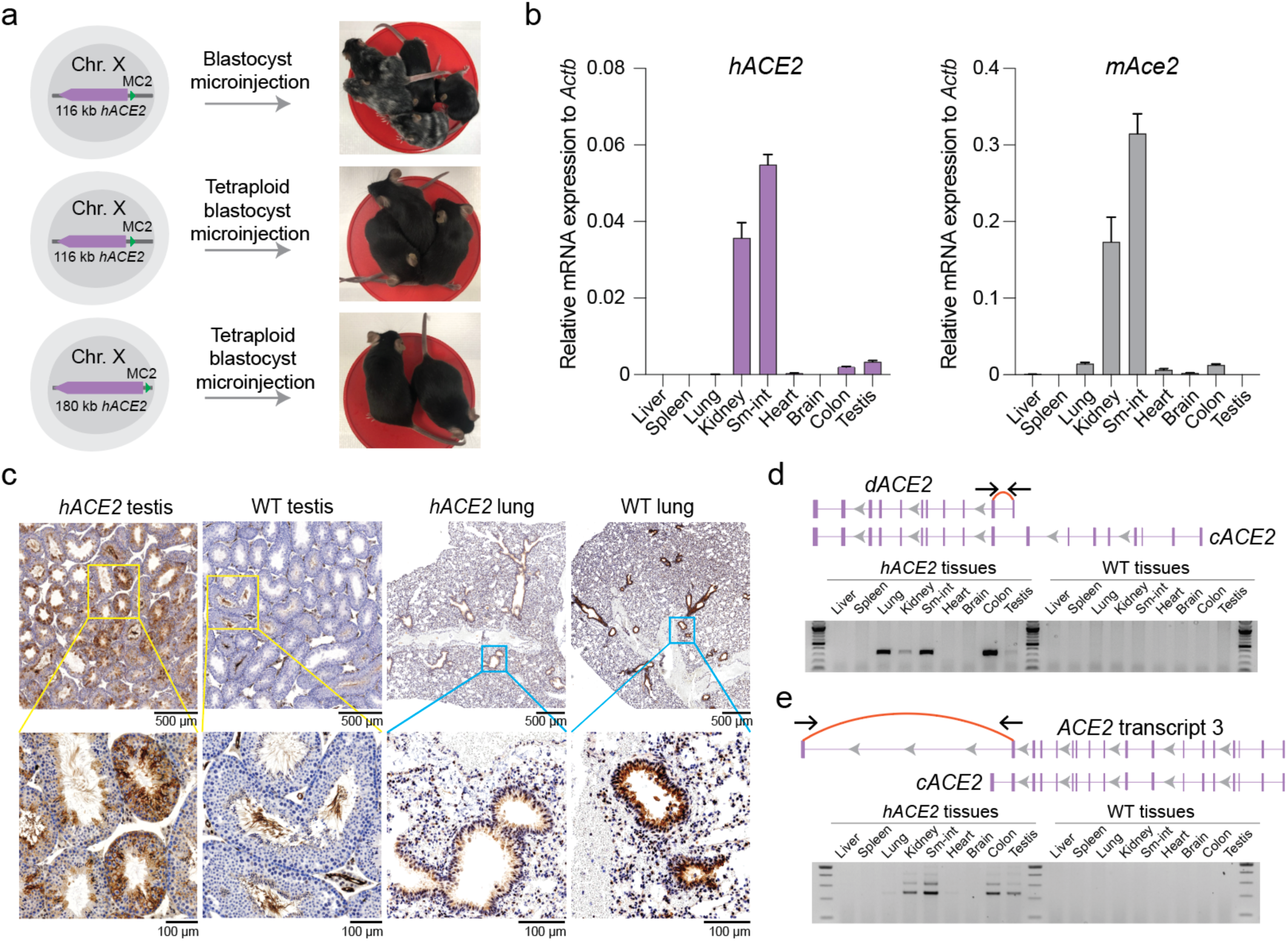
*hACE2* expression characterization. **(a)***hACE2* mice production via chimeric- and tetraploid-blastocyst embryo injection. **(b)** RT-qPCR analysis of *hACE2* (left) and *mAce2* (right) in nine tissues collected from 4-weeks old *hACE2* and wild-type mice. Expression was normalized to mouse *Actb*. Bars represent the mean ± SD of three technical replicates. **(c)** IHC staining analysis of ACE2 in testis and lung dissected from 10-weeks old *hACE2* or wild-type mice. Antibody reacts with both human and mouse ACE2. Yellow and blue boxes mark magnified areas. **(d)** RT-PCR detection of *dACE2* isoform (transcript variant 5) in *hACE2* mouse tissues. *cACE2*, canonical *ACE2* transcript. (**e**) RT-PCR detection of *hACE2* transcript 3 in *hACE2* mouse tissues. *cACE2*, canonical *ACE2* transcript.

Proper spatial expression of *ACE2* is crucial for studying SARS-CoV-2 pathogenesis in mice. Hence, we wondered whether the *hACE2* expression pattern would be recapitulated in the humanized mice. Since the 180 kb *hACE2* payload includes longer *hACE2* upstream sequences including the entire *CLTRN* gene, which may or may not be necessary for the spatiotemporal regulation of *hACE2*, but complicates the humanization scheme by introducing the *hCLTRN* gene; the longer sequences were included as a “backup plan” in case the 116 kb *hACE2* construct was somehow inefficiently expressed. We first examined *hACE2* mRNA expression across nine tissues from the 116 kb *hACE2* GEMM (**Fig. 4b**). Abundant *hACE2* mRNA was detected in small intestine and kidney, while moderate levels were observed in testis and colon, indicating the mouse transcription machinery faithfully expressed *hACE2*. Overall, expression patterns between *mAce2* and *hACE2* were similar aside from a few important differences. For instance, we readily detected *hACE2* in the testis, recapitulating the *ACE2* expression observed in humans, whereas *mAce2* is not expressed in testis of wild-type mice (**Fig. 4b, Fig. S5c**). Importantly, all published humanized *ACE2* models so far failed to express *ACE2* in the testis^24, 36, 37, 60, 61^, making the h*ACE2* mice a valuable resource for modeling possible human testicular infection^62^. In addition, we observed lower *hACE2* expression in the lung of the *hACE2* mice compared with *mAce2* in wild-type mice, consistent with the comparison between human RNA-seq and mouse ENCODE data (**Fig. S5c**). Immunohistochemistry (IHC) staining of *hACE2* testes showed robust ACE2 expression in Sertoli cells, spermatogonia and spermatocytes, reminiscent of the ACE2 expression pattern in human testis^62, 63^. In contrast, only a subset of spermatozoal cells expressed ACE2 in the wild-type testis (**Fig. 4c, Fig. S5d**). IHC staining of lungs showed *ACE2* expression in bronchioles of both *hACE2* and wild-type mice, differed in a much lower level observed in *hACE2* lung (**Fig. 4c**). These data suggest the *hACE2* mice exhibit authentic human tissue-specific gene expression patterns, including some that are missing in non-humanized animals but observed in humans.

Given that we swapped-in the entire *hACE2* gene, we wonder whether human-specific splicing patterns would be recapitulated in the *hACE2* mice. A recent study identified a novel *ACE2* isoform (*dACE2*) as an interferon-stimulated gene, although the product of this transcript is not the receptor of SARS-CoV-2, hinting at a potentially important role of alternative *hACE2* splicing^39, 64^. In our *hACE2* mice, we readily detected *dACE2* in the lung, kidney, small intestine, and colon (**Fig. 4d, Fig. S5e**). In addition, the long *ACE2* transcript (transcript variant 3, **Fig. 3a**) was detected in the small intestine, kidney, brain and testis of *hACE2* mice (**Fig. 4e, Fig. S5f**), further demonstrating that physiological alternative splicing patterns of human *ACE2* are recapitulated in the *hACE2* mice.

### *hACE2* mice are susceptible to SARS-CoV-2 infection

To characterize the susceptibility of *hACE2* mice to SARS-CoV-2, we intranasally challenged the *hACE2, K18-hACE2* and wild-type mice with 10^3^ or 10^5^ plaque-forming units (PFU) of SARS-CoV-2. Given the physiological expression level of *ACE2* in *hACE2* mice (**Fig. 4**), we expected a milder infection manifestation compared to the *K18*-*hACE2* transgene model. All mice were sacrificed on day 3 post-infection (dpi), and viral RNA level in dissected lungs was evaluated by RT-qPCR. As expected, SARS-CoV-2 RNA was undetectable in wild-type lungs; while high levels of SARS-CoV-2 RNA were detected in *K18*-*hACE2* lungs, and these levels positively correlated with inoculum dosage (**Fig. 5a**). For the *hACE2* mice, we detected moderate viral RNA levels in the 10^5^ PFU infection group, and very low amounts in the male *hACE2* mouse of the 10^3^ PFU infection group. Infectious viruses from lung homogenates were quantified using a plaque assay (**Fig. 5b**), and levels were consistent with the RT-qPCR result. We noticed that higher viral RNA levels were detected in male *K18-ACE2* and *hACE2* mouse lungs compared with females, despite identical inoculum dosage. No significant difference in *hACE2* expression was observed between males and females. Notably, *hACE2* mice display ∼70-fold lower *hACE2* expression in lungs compared to transgenic *K18-ACE2* mice (**Fig. 5c**), representing a more physiological expression level. Host interferon stimulated genes *Isg15*, *Cxcl11* and *Mx1* were significantly induced in the *K18-ACE2* mice, and these were moderately induced in the *hACE2* mice, mirroring viral levels (**Fig. S6a**). Transcriptional evaluation of SARS-CoV-2 infected lungs revealed a moderate type I/III interferon response in the *hACE2* mice (**Fig. 5d**), in which the induced genes largely overlap with that of *K18-ACE2* mice, but not with that of wild-type mice (**Fig. 5e, Fig. S6b, Supplementary file 2**). Histopathological examination of infected lung sections revealed that both *K18-ACE2* and *hACE2* mice developed pneumonia evidenced by monocyte infiltration, but *hACE2* mice displayed substantially milder lesions of alveolar epithelial cells (**Fig. 5f**). Corresponding IHC staining showed strong SARS-CoV-2 nucleocapsid protein surrounding the alveolar cells in both models (**Fig. S6c**).

**Fig. 5.**
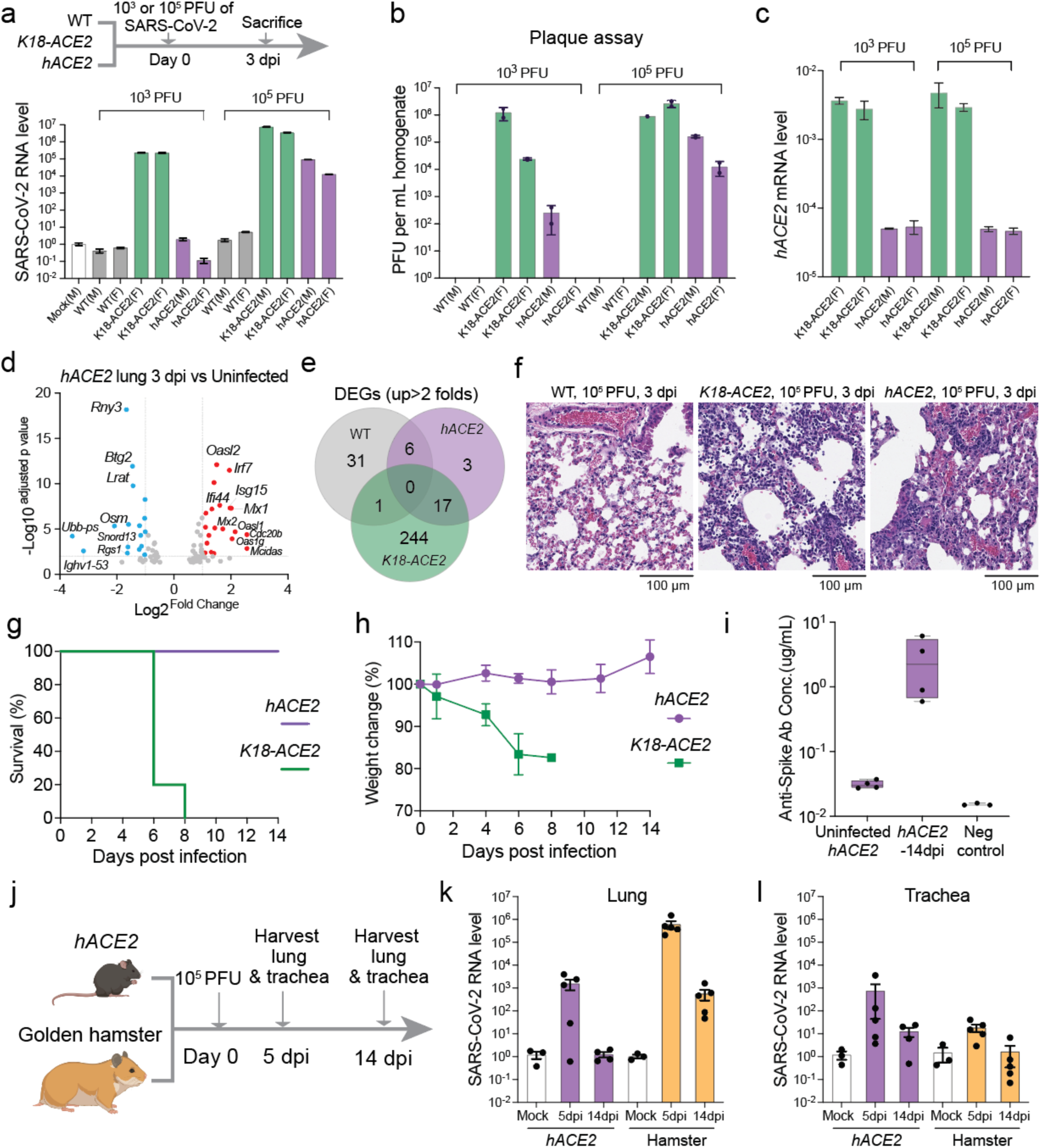
Characterizing the *hACE2* GEMM with SARS-CoV-2 infection. **(a-c)** Lungs dissected from wild-type (WT), *K18-ACE2* and *hACE2* mice infected with SARS-CoV-2 were analyzed for SARS-CoV-2 nucleocapsid gene expression by RT-qPCR (a), infectious viral levels by plaque assay (b), and human *ACE2* expression by RT-qPCR (c). SARS-CoV-2 levels were normalized *Actb* and to an uninfected control. Human *ACE2* expression levels were normalized *Actb*. Bars represent mean ± SD of three technical replicates. M, male mice; F, female mice. **(d)** Volcano plot of *hACE2* infected lungs vs uninfected lungs. Red colored genes are upregulated and blue colored genes are downregulated in infected lungs. Fold change cutoff is set at 2, adjusted p-value cutoff is set at 0.01. **(e)** Venn diagram of upregulated (cutoff is 2-fold) differentially expressed genes (DEGs) in wild-type, *K18-ACE2* and *hACE2* lungs. **(f)** Histopathological analysis of wild-type, *K18-ACE2* and *hACE2* female lungs by hematoxylin and eosin (H&E) staining. (**g-h**) *K18-ACE2* (n=5) and *hACE2* (n=4) mice were intranasally infected with 10^5^ PFU of SARS-CoV-2 and were monitored every other day for morbidity (g) and weight (h). Bars represent mean ± SD of biological replicates. **(i)** Serological detection of anti-SARS-CoV-2 mouse IgG via enzyme-linked immunoassay (ELISA). **(j)** Schematic of the longitudinal SARS-CoV-2 infection experiment for *hACE2* mice and wild-type golden hamsters. Mouse and hamster icons were created with BioRender. (**k-l**) RT-qPCR analysis of SARS-CoV-2 nucleocapsid gene in the lung (k) and trachea (l) of *hACE2* mice and hamsters. *hACE2* mice, n=5 (5 dpi), 4 (14 dpi). Hamsters, n=5 (5 dpi), 5 (14 dpi). SARS-CoV-2 RNA level was normalized *Actb* and to uninfected control. Bars represent mean ± SEM of biological replicates.

Human COVID-19 is a complex disease with very diverse manifestations and outcomes reflecting the age, health status, immune status and genetic makeup of human patients. We thus tested whether the *hACE2* model could be used to better model human SARS-CoV-2 infection as compared to the *K18-hACE2* model, which is known to succumb to SARS-CoV-2 within 10 days^24^, and is thus unable to recapitulate even the medium-term, let alone long-term effects of viral infection. We infected *hACE2* and *K18-ACE2* mice with 10^5^ PFU of SARS-CoV-2, and monitored weight and survival over the course of 14 days. All *hACE2* mice survived to the end without obvious sickness. In contrast, *K18-ACE2* had significantly reduced mobility 5 days post infection; 4 out of the 5 mice died at 6 dpi, and 1 died at 8 dpi (**Fig. 5g**). Body weight measurement showed that *K18-ACE2* mice lost weight substantially prior to fatality, whereas the *hACE2* mice did not (**Fig. 5h, Fig S6d**). Measurements of antiviral humoral immune response by ELISA showed evidence of anti-Spike trimer antibody in the 14 dpi *hACE2* sera (**Fig. 5i**). These data collectively suggest that *hACE2* mice can recover from SARS-CoV-2 infection, and are thus likely to be particularly useful for modeling various aspects of human COVID-19 pathophysiology.

The golden hamster (*Mesocricetus auratus*) is another commonly used rodent model for studying infection with respiratory viruses^65–68^. However, such studies are limited by the lack of genetic tools, and a very limited repertoire of hamster mutants that could be used to model comorbid conditions. We wondered whether the *hACE2* mice were comparable to hamster in terms of susceptibility to SARS-CoV-2. We set up a longitudinal infection experiment, including collection of lungs and tracheas on 5 dpi and 14 dpi (**Fig. 5j**). SARS-CoV-2 viral RNA was detected on 5 dpi in the lung of both *hACE2* mice and hamsters albeit detection was more moderate for *hACE2* mice, and diminished significantly on 14 dpi (**Fig. 5k**). In hamster trachea, viral RNA levels very mildly increased at 5 dpi (**Fig. 5l**), probably due to the lack of *Ace2* expression in hamster’s tracheal epithelial cells^69^. In contrast, higher levels of viral RNA were detected in trachea of a subset of *hACE2* mice (**Fig. 5l**), consistent with previous results in human patients^70, 71^. Taken together, *hACE2* GEMM has a milder but comparable infectibility with golden hamster in lungs, and perhaps a more human-like infectibility in trachea.

## Discussion

Understanding the basis of mammalian genomes is underway from many different perspectives. Advanced genome sequencing technologies have already revealed the complex genetic blueprints of many vertebrates^72, 73^. To directly probe the roles of regulatory components and genome polymorphism we provide here a strategy to reliably overwrite hundreds of kilobases of native mammalian genomic segments with carefully designed synthetic DNAs or cross-species gene counterparts. Mammalian genome writing is ideal for introducing tens to hundreds of edits through *de novo* synthesis, which otherwise is extremely difficult, if not impossible, to engineer with traditional genome editing approaches such as CRISPR, not to mention maintaining cells’ developmental potential through multiple rounds of editing. The iterative genome writing nature of mSwAP-In overcomes the size limitation of DNAs to be delivered, paving the way for eventual writing of megabase-sized synthetic DNAs. The combination of positive selection and counterselection ensures on-target integration of payload DNA. In conjunction with targeted capture sequencing, clones with undesirable genomic outcomes (e.g. integration of plasmid backbones, or co-transfected plasmids, as well as structural payload variants) can be identified and eliminated, reducing experimental bias.

Although we demonstrated that mSwAP-In can be used to deliver both homologous and non-homologous DNA sequences to mESCs, we believe mSwAP-In can be generalized to many other mammalian systems provided that homologous recombination is comparably efficient to mESCs. Further optimizations may increase efficiency and simplify the workflow. For example, 1) Inhibiting the non-homologous end joining pathway during nucleofection; 2) Using exogenous counter-selectable markers in MC2 to circumvent deletion of endogenous *Hprt*; and 3) Engineering a piece of reference DNA (e.g. *Actb*) into the capture sequencing bait to determine the copy number of integrated payload DNA directly from sequencing data.

The mouse is the most frequently used mammalian model in clinical studies. However, recapitulation of human diseases in mice often fails due to evolutionary differences. Genetically humanizing complete mouse loci by *in situ* replacement provides a means to improve disease recapitulation accuracy since human-specific spatiotemporal regulation and splicing are more likely to be preserved. Because of the high efficiency of mSwAP-In, producing a large number of informative GREAT-GEMMs in a short period of time is possible.

In response to the COVID-19 outbreak, we rapidly generated a genomically humanized *ACE2* mouse model with mSwAP-In. In contrast to existing humanized *ACE2* models, we found that the *ACE2* expression level and distribution more closely resembled that in humans (**Fig. 4**). In terms of susceptibility of the *hACE2* mice to SARS-CoV-2, we found they are readily infected, but display mild disease symptoms without mortality, and showed evidence of a humoral antiviral response, similar to the outcomes observed in most healthy younger humans. We think this *hACE2* model is a valuable platform for studying the long-term effects of COVID-19 *in vivo*. In addition, mortality and more severe symptoms are more common in elderly individuals and people with comorbidities. The *hACE2* mice used in this study were relatively young (10-15 weeks old) and healthy, corresponding to young people with mild or minimal COVID-19 symptoms. Infection experiments using older *hACE2* mice or combining *hACE2* with existing mouse models, such as diabetes, obesity etc., may be informative for modeling severe COVID-19. Finally, we showed that *hACE2* mice behaved similarly upon infection compared to golden hamsters, a commonly used rodent model, which lacks adequate genetic resources for thorough modeling of COVID-19.

## Materials and Methods

### BACs plasmids

Human (CH17-203N23, CH17-449P15) and mouse (RP23-51O13, RP23-75P20) BACs were purchased from BACPAC Resources Center. Yeast-bacterium shuttle vector pLM1050 was modified by Dr. Leslie Mitchell based on a previous study^16^. pWZ699 was constructed by inserting a cassette containing pPGK-Λ1TK-SV40pA transcription unit and the *Actb* gene into the *Not*I site of pLM1050. Marker cassette 1 donor plasmids for *synTrp53* and *hACE2* loci were constructed using Gibson assembly of MC1 and two homology arms into pUC19 vector. ∼ 750 bp left and right homology arms were amplified from BACs. When using microhomology-mediated end joining for MC1 insertion, 20 bp micro-homology arms were carried on primers. pX330 plasmid was purchased from Addgene (42230).

### Mammalian cell lines, yeast strain

The C57BL/6J mESC line (MK6) was obtained from NYU Langone Health Rodent Genetic Engineering Core. Both feeder-dependent and feeder-independent culture conditions were used for different purposes in this study. The medium for feeder-dependent condition consists of 85% (v/v) KnockOut DMEM (Fisher Scientific, 10829018), 15% (v/v) Fetal Bovine Serum (Hyclone, SH30070.03), 0.5 mg/ml Penicillin-Streptomycin-Glutamine (Gibco, 10378016), 7 μL 2-Mercaptoethanol (Sigma-Aldrich, M6250), 0.1 mM MEM Non-Essential Amino Acids (Gibco, 11140050) and 1 U/ml LIF (EMD Millipore, ESG1107). Tissue culture treated plates were first coated with 0.1% gelatin solution (EMD Millipore, ES-006-B), followed by seeding 5×10^4^/cm^2^ of mouse embryonic fibroblast (MEF) cells (CellBiolabs, CBA-310) in MEF medium (DMEM [Gibco, 11965118], 10% Fetal Bovine Serum [GeminiBio, 100-500], 0.1 mM MEM Non-Essential Amino Acids, 2 mM L-glutamine, 1% Pen-Strep). mESCs were plated on the MEF monolayer. Feeder-independent medium consisted of 80% of 2i basal medium supplement with 3 µM CHIR99021 and 1 µM PD0325901, 20% of feeder-dependent ES medium (mentioned above). Tissue culture treated plates were coated with 0.1% gelatin solution before use. All cells were grown in a humidified tissue culture incubator at 37°C supplied with 5% CO_2_. VeroE6 cells (kidney epithelial cells from female African green monkey, ATCC, CRL-1586) were cultured in 12-well plates with DMEM supplemented with 4% FBS, 1% penicillin-streptomycin-neomycin (PSN), and 0.2% agarose (Lonza, 50100). BY4741 yeast strain was used for all the payload assemblies.

### Virus

SARS-CoV-2 strain USA-WA1/2020 (NR-52281) was obtained from BEI Resources, NIAID, NIH (Bethesda, MD, USA). SARS-CoV-2 viruses were expanded in VeroE6 cells^65^. Harvested viruses were purified with Amicon Ultra-15 Centrifugal filter unit (Millipore Sigma). The SARS-CoV-2 virus stock titer was determined by performing a plaque assay in VeroE6 cells.

### Animals

Engineered mESCs were either injected into C57BL/6J-albino (Charles River laboratories, strain#493) blastocysts, or injected into B6D2F1/J (Jackson laboratories, strain#100006) tetraploid blastocysts for mice production. Mice were housed in NYU Langone Health BSL1 barrier facility. Wild-type C57BL/6J (strain#000664) and *K18*-*hACE2* (strain#034860) mice were obtained from The Jackson laboratory. Golden hamsters were obtained from Charles River Laboratories (strain#049). Ten to fifteen weeks old mice and ten to twelve weeks old hamsters were transferred to the NYU Langone Health BSL3 facility for the SARS-CoV-2 infection. All mice were settled for at least two days prior to infection. All experimental procedures were approved by the Institutional Animal Care and Use Committee (IACUC) at NYU Langone Health.

### Payload DNA assembly and preparation

Two approaches were used for payload DNA assembly in this study. For synthetic *Trp53* and its subsequent 40 kb, 75 kb and 115 kb payloads, DNA fragments ranging from 3 kb to 5 kb with 40-100 bp terminal homologies were amplified from mouse BAC RP23-51O13 with Q5 polymerase (NEB, M0491L). Approximately equal amount (100 ng) of each PCR fragment, together with 50 ng of each linker fragment for bridging vector and insert and 20 ng linearized pLM1050 vector were co-transformed into yeast for assembly. For *hACE2* payloads, CH17-203N23 and CH17-449P15 BACs were extracted by using a NucleoBond Xtra BAC kit (Takara, 740436.25). ∼1 μg of BAC DNA was digested with 30 nM of sgRNAs (IDT), and 30 nM recombinant Cas9 nuclease (NEB, M0386S) at 37°C for 2 hours. 1 μL of 20 mg/ml proteinase K was added to the digestion reaction for 10 minutes at room temperature. Digested BAC and *Sal*I-linearized acceptor vector were co-transformed into yeast for assembly. Yeast cells were cultured on SC–Leu plates at 30°C for 3 days. Yeast colony containing correct payload was identified by screening all novel junctions between each two fragments. To assemble the 180 kb *hACE2* payload, an *URA3* gene was inserted in front of the MC2 of the 116 kb *ACE2* payload. The 64 kb *ACE2* region of interest was released from CH17-449P15 BAC by *in vitro* Cas9-gRNA digestion. A plasmid expresses Cas9 and gRNA targeting *URA3* in yeast was co-transformed with the 64 kb *ACE2* fragment into BY4741 strain. Yeast cells were selected with 5-FOA for successful insertion of the 64 kb *hACE2* fragment. Payload DNA was isolated from yeast by using a yeast plasmid miniprep kit (Zymo Research, D2001), eluted in 30 μL of TE. 2 μL of yeast miniprep DNA was used for electroporation into EPI300 *E. coli* strain (Lucigen, EC300150). *E. coli* colonies containing payload DNAs were grown in a 5 ml LB plus 50 μg/mL kanamycin culture overnight, and diluted at 1:100 ratio into 250 ml LB supplemented with kanamycin (50 μg/mL) and 1x copy number induction solution (Lucigen, CCIS125). Payload DNA was isolated from *E. coli* by using a NucleoBond Xtra BAC kit (Takara, 740436.25) for delivery into mESCs. Primers used for assembly are listed in Supplementary file 3.

### BAC and payload DNA sequencing library construction

Concentration for BACs and assembled payload DNAs was quantified by using a Qubit dsDNA HS kit (Thermo Fisher, Q32854), Approximately 100 ng DNA was used for the library construction using the NEBNext Ultra II FS DNA library prep kit (E7805). AMPure XP beads (Beckman Coulter, A63881) were used for DNA purification on a magnetic stand. DNA libraries were loaded on a ZAG DNA analyzer (Agilent) for quality control. DNA libraries were sequenced on an Illumina NextSeq 500.

### Sequencing data processing

Sequencing reads were demultiplexed using bcl2fastq v2.20, and subsequently trimmed using Trimmomatic v0.39. Trimmed reads were aligned to references using BWA v0.7.17. Duplicates were marked using samblaster v0.1.24. Coverage depth tracks and quantification was generated using BEDOPS v2.4.35. Sequencing data were visualized using UCSC genome browser.

### Pulse-field gel electrophoresis

Payload DNAs were linearized using a single-cut restriction enzyme, followed by heat inactivation as recommended by the manufacturer. 200 ng of digestion product was loaded into a 1% low melting point agarose gel. Lambda-PFG ladder (NEB, N0341S) or lambda DNA-Mono cut mix (NEB, N3019S) were used as ladders. CHEF Mapper XA System (Bio-Rad), auto-algorithm was used for electrophoresis. Agarose gel was first stained with 0.5 μg/mL ethidium bromide in deionized water for 30 min, and then destained with deionized water for 30 min before imaging on a ChemiDoc MP imaging system (Bio-Rad).

### Crystal violet staining

mES clones grown on gelatin coated plate were washed with PBS once, then fixed in 4% (w/v) formaldehyde for 15 minutes at room temperature followed by two rounds of washing with PBS. 0.1% (diluted with 10% ethanol) crystal violet (Sigma-Aldrich, V5265) dye was used to stain the mES colonies for 20 minutes at room temperature followed by three rounds of washing with water. Plates were air dried at room temperature before counting the colony number.

### Nucleofection

Depending on the culture conditions, 10 cm tissue culture dishes were pre-coated with either 0.1% gelatin (EMD Milipore, ES-006-B) or 2×10^6^ mitomycin treated MEF feeder cells. mESCs were trypsinized with 0.25% Trypsin-EDTA (Gibco, 25200056) at 37°C for 6 minutes. Cell number was determined by hemocytometer. Approximately 3 million mESCs were washed with DPBS (Gibco, 14190144) and pelleted by centrifugation at 300 x g for 5 minutes at room temperature. A total of 10 μg DNA mixture containing payload DNA and Cas9-gRNA plasmid(s) (Table S3) was used for the nucleofection. Nucleofection solutions and cuvette were from Mouse ES Cell Nucleofector kit (Lonza, VPH-1001). Nucleofector (Lonza 2b) A-023 program was used to deliver the DNA mixture into mESCs. Nucleofected mESCs were plated onto pre-coated 10 cm dishes, and cultured in 37°C, 5% CO_2_ humidified incubator.

### mESCs colony picking and PCR screening

Mitotically inactivated MEFs were pre-seeded in a 96-well tissue culture plate (Corning, 3595) in MEF medium one day before colony picking. The next day, MEF medium was changed to 100 μL/well of ES medium at least 2 hours before use. 10 cm plates containing mES colonies were wash with PBS once, and refilled with 10 mL of PBS. mES colonies were aspirated with 10 μL of PBS using a P20 pipet, and transferred to an empty round bottom low retention 96 well plate (Corning, 7007). 35 μL/well of accutase (Gibco, A1110501) was added to the mES colonies for dissociation at 37°C for 9 min. 100 μL/well of ES medium was used to neutralize the trypsinization. mESCs were singularized by at least 20 times of gentle pipetting. 100 μL of the cell suspension was transferred to a gelatin-coated 96 well plate prefilled with 100 μL of ES medium. The rest of cell suspension (∼40 μL) was transferred to the 96 well MEF plate prefilled with 100 μL of ES medium. ES medium was refreshed daily until the feeder-independent plate becomes >50% confluent. mESCs from feeder-independent plate were trypsinized and 10% cells were passaged to a new gelatin-coated plate for proliferation, 90% of cells were transferred to a PCR plate. mESCs in the PCR plate were span down at 300 x g for 5 min, and supernatant was discarded. Cell pellets were resuspended with 30 μL of lysis buffer (0.3 μg/ml proteinase K in TE). mESCs were lysed on a thermal cycler using 37°C 1 hour, 98°C 10min, 16°C keep program. 1 μL of mESC lysate was used as template in a 10 μL PCR reaction.

### Digital PCR for human *ACE2* copy number determination

Genomic DNA of mESCs was extracted by using a QIAamp DNA mini kit (QIAGEN, 51306). For *hACE2* mESCs, approximately 500 ng of gDNA and *hACE2* payload DNA containing *mActb* gene on the backbone were digested with *Eco*RI (NEB, R3101S) at 37°C for 2 hours. 50 ng digested mESC gDNA and 1 pg digested *hACE2* payload DNA were used for qPCR analysis. For *synTrp53* mESCs, a wild-type mESC gDNA sample digested with *Eco*RI in the sample way as candidates, was used as normalization control. SYBR Green Master Mix (Roche, 04887352001) was used for the qPCR reaction on a LightCycler 480 instrument. Copy number was normalized to *hACE2-mActb* payload (for *hACE2* clones) or wild-type mESCs (for *synTrp53* clones).

### mESCs capture sequencing library construction

1-3 million of feeder-independent mESCs were harvested for genomic DNA extraction by using a QIAamp DNA Mini Kit (QIAGEN, 51306). Genomic DNA concentration was determined with a Nanodrop spectrophotometer, ∼1 μg genomic DNA was subjected to DNA library construction using a large fragment size kit (NEBNext Ultra II FS). Final DNA library concentration was measured using a Qubit dsDNA HS assay kit (Invitrogen, Q32851). For *synTrp53* mESCs, capture bait comprises RP23-51O13, marker cassette 1, marker cassette 2 and pX330 (addgene, 42230). For *hACE2* humanized mESCs, capture bait comprises CH17-203N23, CH17-449P15, RP23-75P20, marker cassette 1, marker cassette 2 and pX330. Bait DNA mixture was labeled with Biotin-16-dUTP (Roche, 11431692103) using a nick translation kit (Sigma-Aldrich, 10976776001). The capture was performed as previously described^14^. Briefly, biotinylated baits DNA mixture was prehybridized, and mix with DNA library samples at 65°C for 16 to 22 hours. Captured DNA was purified using Streptavidin C1 beads (Invitrogen, 65002) and amplified using KAPA Hi-Fi Hotstart PCR kit (Roche, KK2602). After one more step of DNA cleanup, capture library was sequenced on an Illumina NextSeq 500 using a 75 cycles kit.

### *mAce2* and *hACE2* mRNA RT-qPCR

Mouse tissues were dissected and homogenized using a pellet pestle (Fisher Scientific, 12141364). Total RNA was extracted using a RNeasy kit following vendor’s instructions (QIAGEN, 74136). Approximately 1 μg of total RNA was used for reverse transcription (Invitrogen, 18091050). 1 μL of 1:10 diluted cDNA was used in a 10 μL SYBR Green (Roche, 04887352001) qPCR reaction on a LightCycler 480 instrument (Roche). Primers used for RT-qPCR are listed in Table S4.

### *In vivo* SARS-CoV-2 infection

C57BL/6J, *K18-hACE2* and *hACE2* mice (this study) were anesthetized with intraperitoneal injection of 150 μL ketamine (10 mg/mL)/xylazine (1 mg/mL) solution. Hamsters were injected with 200 μL of ketamine (75 mg/mL)/xylazine (5 mg/mL in PBS) solution. 1×10^3^ or 1×10^5^ PFU of SARS-CoV-2 were administered intranasally in a total volume of 50 μL PBS per mouse, 100 μL PBS per hamster, delivered to both nostrils equally. All infection experiments were performed in the NYU BSL3 facility.

### SARS-CoV-2 infected lung and trachea RNA extraction and quantification

One lobe of lung was immersed in 1 ml Trizol solution (Invitrogen, 15596018) in Lysing Matrix A homogenization tubes (MP Biomedicals) immediately after dissecting from euthanized mouse or hamster. Lung was homogenized following manufacturer’s instructions (MP Biomedicals, FastPrep-24 5G). Trachea was dissected and immersed in 1 mL PBS in a 2 mL microcentrifuge tube (Fisherbrand, 14-666-315) containing one stainless steel bead (QIAGEN, 69989). After the homogenization, PBS homogenates were centrifuged for 2 minutes at 5,000 x g. 500 μL of homogenates were transferred and mixed with 500 μL Trizol solution for RNA extraction. Processing lung and trachea sample for the following steps, 200 μL of chloroform per 1 ml of Trizol reagent was added and vortexed thoroughly. Tubes were centrifuged at 12,000 x g for 10 min at 4°C. Aqueous phase was transferred to a new RNase-free 1.5 mL tube. Total RNA was precipitated by adding 500 μL of isopropanol per 1 ml Trizol solution, and pelleted by centrifugation at 12,000 x g for 10 min at 4°C. RNA pellet was washed with 500 μL of 75% ethanol once, air-dried at room temperature for 10 min, and dissolved with 100 μL of RNase-free water. Total RNA from SARS-CoV-2 infected lung or trachea was subjected to one-step real-time reverse transcription PCR using One-step PrimeScript RT-PCR kit (Takara, RR064B). Multiplex PCR was performed to detect SARS-CoV-2 nucleocapsid gene and mouse *Actb* gene. Probe targeting SARS-CoV-2 was labeled with FAM fluorophore and probes targeting *Actb* gene was labeled with Cy5 fluorophore. RT-PCR was performed on a LightCycler 480 instrument. SARS-CoV-2 RNA level was normalized to *Actb*.

### Lung RNA sequencing and analysis

Lung total RNA quality and quantity were examined using a Bioanalyzer (Agilent 2100, RNA 6000 nano kit). Sequencing libraries were constructed using a TruSeq Stranded Total RNA Library Prep Gold kit (Illumina, 20020599). Libraries were sequenced on an Illumina NovaSeq 6000 using a SP100 reagent kit (v1.5, 100 cycles). RNA-seq data were analyzed by using the sns rna-star pipeline. Briefly, adapters and low-quality bases were trimmed using Trimmomatic (v0.36). Sequencing reads were mapped to the mouse reference genome (mm10) using the STAR aligner (v2.7.3). Alignments were guided by a Gene Transfer Format (GTF) file. The mean read insert sizes and their standard deviations were calculated using Picard tools (v.2.18.20). The genes-samples counts matrix was generated using featureCounts (v1.6.3), normalized based on their library size factors using DEseq2, and differential expression analysis was performed. The Read Per Million (RPM) normalized BigWig files were generated using deepTools (v.3.1.0). Data were visualized using GraphPad Prism or Rstudio.

### Plaque assay

The second lobe of lung or trachea was immersed in 1 mL PBS in a 2 mL Micro Centrifuge Tube (Fisherbrand, 14-666-315) containing one stainless steel bead (5 mm, QIAGEN, 1026563) immediately after dissecting the SARS-CoV-2 infected mouse or hamster. Lung or trachea was homogenized following manufacturer’s instructions (TissueLyser II, QIAGEN, 85300). Homogenates were then centrifuged for 2 minutes at 5,000 x g and immediately frozen until plaque assay was performed. Plaque assay was performed with VeroE6 cells (ATCC, CRL-1586) plated in 24-well plates. Samples were diluted logarithmically in Minimal Essential Media (Gibco, 11095072), of which 200 μL were inoculated per well and incubated for 1 hour at 37°C. Inoculated cells were then overlayed with DMEM supplemented with 4% FBS, 1% penicillin-streptomycin-neomycin (PSN), and 0.2% agarose (Lonza, 50100). Overlayed cells were incubated at 37°C for 48 hours and subsequently fixed with 10% neutral buffered formalin for 24 hours. Remaining VeroE6 cells were stained with 0.2% crystal violet in 20% ethanol for 10 minutes.

### Histology

The accessary lung lobes were immersed in 5 ml of 10% formalin solution (Sigma-Aldrich, HT501128) for 24 hours at room temperature, and processed through graded ethanols, xylene and paraffin in a Leica Peloris automated processor. Five-micron paraffin-embedded sections were either stained with hematoxylin (Leica, 3801575) and eosin (Leica, 3801619) on a Leica ST5020 automated histochemical stainer or immunostained on a Leica BondRX® autostainer, according to the manufacturers’ instructions. In brief, sections for immunostaining underwent epitope retrieval for 20 minutes at 100°C with Leica Biosystems ER2 solution (pH 9.0, AR9640). Sections were incubated with one of the two ACE2 antibodies (Thermo, MA5-32307, clone SN0754 or Abcam, ab108209, clone EPR4436) diluted 1:100 for 30 minutes at room temperature and detected with the anti-rabbit HRP-conjugated polymer and DAB in the Leica BOND Polymer Refine Detection System (DS9800). Alternatively, sections were blocked with Rodent Block (Biocare, RBM961 L) prior to a 60 min incubation with anti-SARS-CoV-2 N (Thermo, MA1-70404, clone B46F) diluted 1:100 and then a 10 min incubation with a mouse-on-mouse HRP-conjugated polymer (Biocare MM620 H) and DAB (3,3′-Diaminobenzidine). Sections were counter-stained with hematoxylin and scanned on either a Leica AT2 or Hamamatsu Nanozoomer HT whole slide scanner.

### ELISA

Mouse blood was collected via cardiac puncture, and isolated serum was diluted 100-fold using the dilution buffer of a mouse anti-SARS-CoV-2 antibody IgG titer serologic assay kit (ACROBiosystems, RAS-T023). Diluted samples were added to a microplate with pre-coated SARS-CoV-2 Spike protein (2 μg/mL), and incubated at 37°C for 1 hour. Following 3 washes, 100 μL of HRP-goat anti-mouse IgG (80 ng/mL) was added to the microplate and incubated at 37°C for 1 hour. Following another 3 washes, 100 μL of substrate solution was added and incubated 37°C for 20 min. The reaction was stopped by adding 50 μL stop solution, the absorbance was measured at 450 nm and 630 nm using an imaging reader (BioTek, Cytation 5). Absorbance values for the serum samples were calculated by subtracting the value of the A_630nm_ from the value at A_450 nm_. A standard curve was generated using a series of diluted anti-SARS-CoV-2 mouse IgG control samples. Anti-SARS-CoV-2 mouse IgG titer in mouse serum was quantified using a standard curve.

## Supporting information

Bamintersect analysis

DEGs list

Primer sequences

## Acknowledgements

We thank Francisco Sánchez-Rivera for suggesting the name GREAT-GEMMs. We thank the NIH/NHGRI for supporting this work via CEGS grant 1RM1HG009491. We thank Ludovic Desvignes for access to and help with the NYU BSL3 facility. We thank members of the Experimental Pathology Research Laboratory, which is partially supported by the Cancer Center Support Grant P30CA016087 at NYU Langone’s Laura and Isaac Perlmutter Cancer Center. We thank Ziyan Lin at the Applied Bioinformatics Laboratories NYU Langone Health for the swift processing of RNA sequencing data.

## Author contributions

W.Z. and J.D.B. conceptualized the study. W.Z., I.G., B.R.t., and J.D.B. designed the experiments. W.Z. constructed the payload constructs in this study and delivered into mESCs. W.Z., S.Y.K., A.M.W. and Y.Z. established the *hACE2* breeding colonies. R.B., Y.Z., E.H., H.A. and M.T.M. performed capture sequencing and data analysis. I.G., L.C. performed the SARS-CoV-2 infection and mouse tissue harvesting in the BSL3 facility. P.D.Y., C.K., K.M.K. assisted with mouse experiments in the BSL3 facility at NYULH. W.Z. wrote the manuscript. J.D.B., B.R.t. and R.B. reviewed and edited the manuscript.

## Extended data

**Fig. S1.**
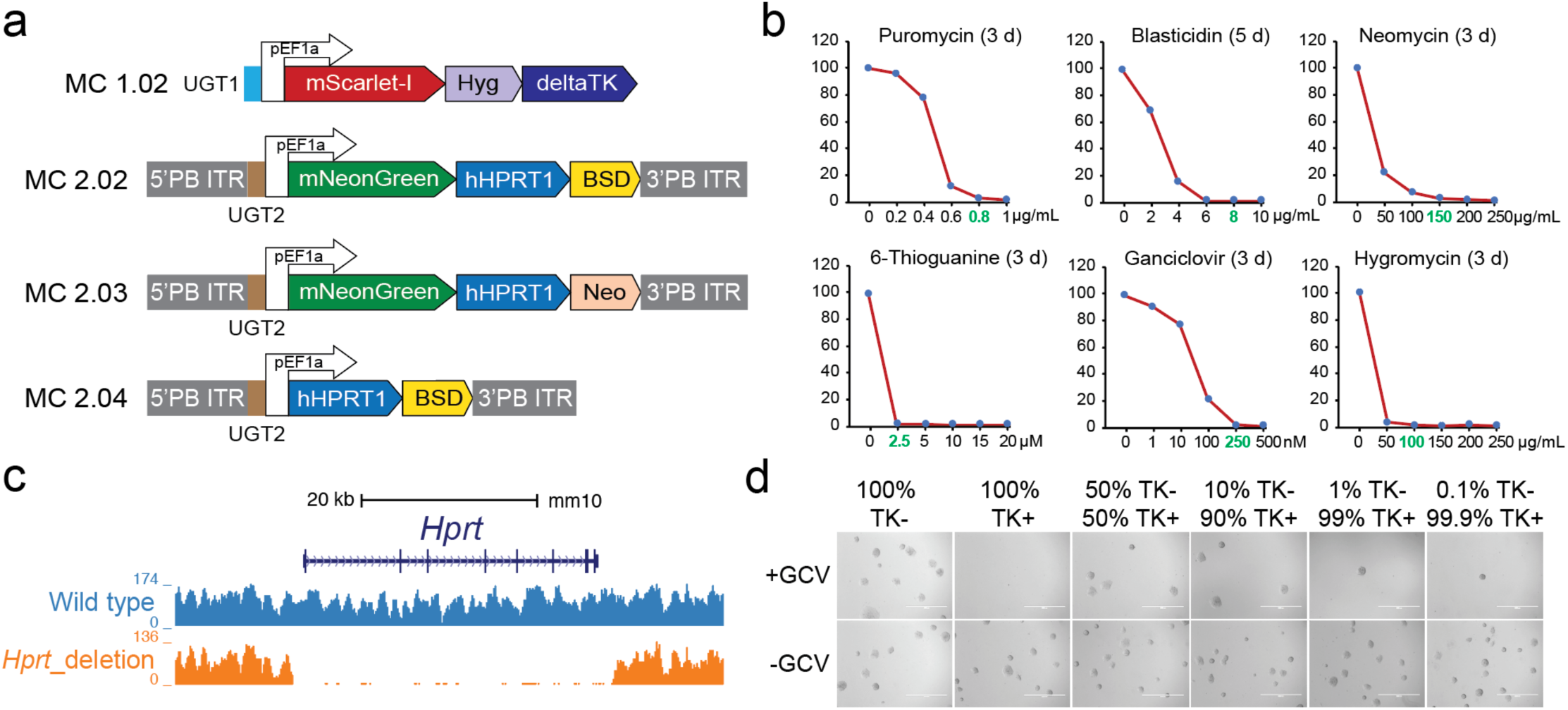
(*match to* Figure 1) mSwAP-In design and development. (**a**) Alternative marker cassettes compatible with genetic backgrounds harboring preexisting drug resistance genes. PB ITR, piggyBac inverted terminal repeat; UGT, universal gRNA target. (**b**) mESC kill curve for each mSwAP-In selection marker. Selected concentrations are highlighted in green: 0.8 μg/ml for puromycin, 8 μg /ml for blasticidin, 150 μg/ml for neomycin, 2.5 μM for 6-thioguanine, 250 nM for ganciclovir and 100 μg/ml for hygromycin. (**c**) Capture-seq analysis of *Hprt* deletion. Sequencing reads were mapped to mm10. (**d**) The bystander effect of thymidine kinase can be overcome by plating single colonies. As few as 0.1% TK-negative cells can be isolated. GCV, ganciclovir (250 nM).

**Fig. S2.**
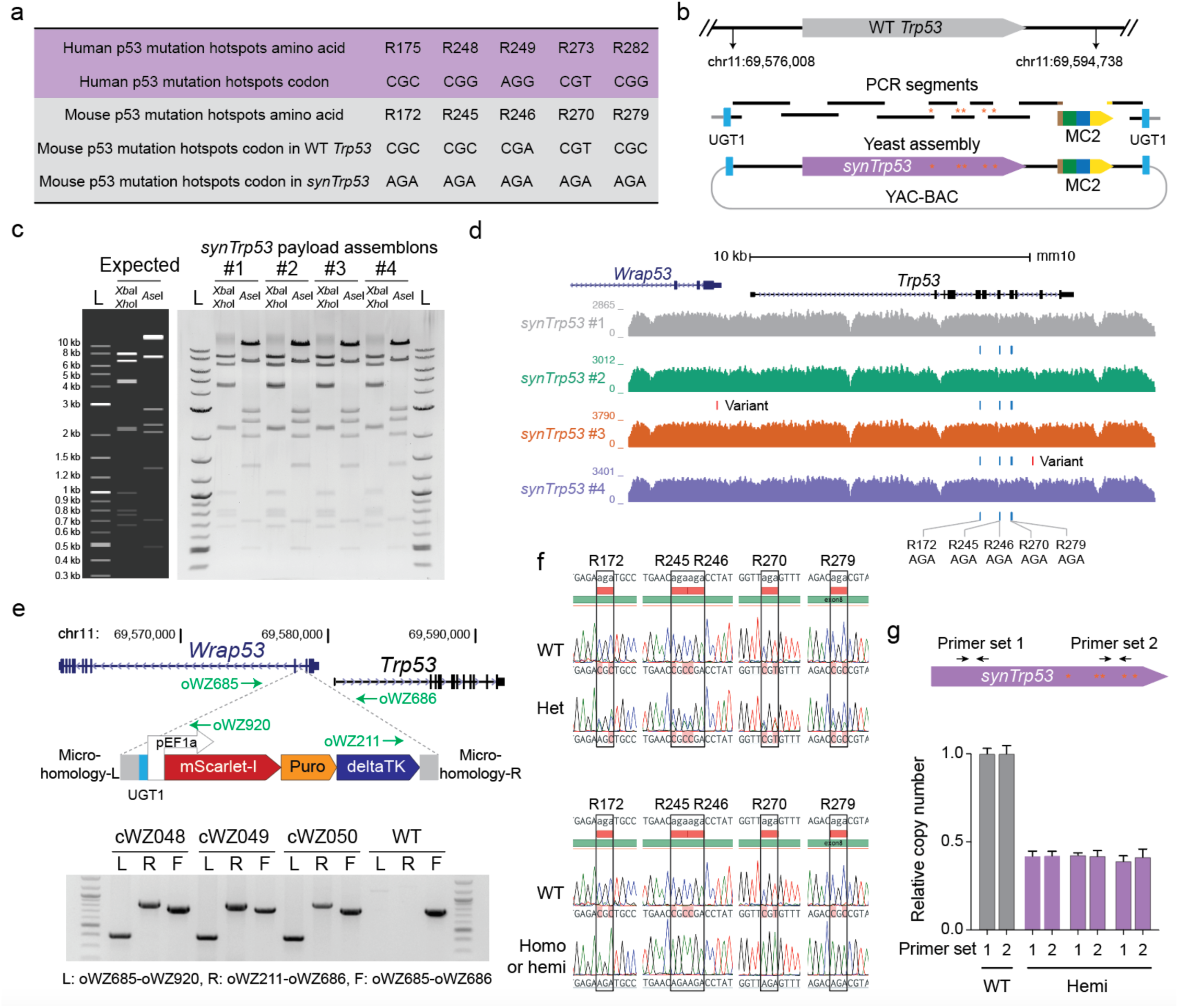
(*match to* Figure 2) Synthetic *Trp53* mSwAP-In. (**a**) p53 mutation hotspots and the corresponding DNA codons in human and mouse, as well as the recoded codons in *synTrp53*. (**b**) *SynTrp53* assembly workflow. Red asterisks represent the recoded codons. (**c**) Restriction enzyme digestion verification of *synTrp53* assemblons. *Xba*I+*Xho*I and *Ase*I were used for each candidate. Predicted digestion patterns were simulated using Snapgene software. L, 1 kb plus ladder (NEB). (**d**) Sequencing coverage of *synTrp53* payload candidates. Reads were mapped to mm10 reference. Clones 1 and 3 have expected variants reflecting the recoded codons in synthetic *Trp53*. Clone 2 and clone 4 have additional undesired variants likely introduced by PCR. (**e**) Marker cassette 1 insertion into the second intron of *Wrap53*. 20 bp microhomology arms were added to each end of MC1 during plasmid construction. Successful insertion was verified by junction PCR. Primers are indicated as green arrows. (**f**) Sanger sequencing validation of heterozygous and hemizygous *synTrp53* integrants. **(g)** *Trp53* copy number qPCR analysis for the three mESC clones only carrying recoded codons (Fig. 2b). *Trp53* copy number was normalized to *Pgk1* gene. Bars represent mean ± SD of three technical replicates.

**Fig. S3.**
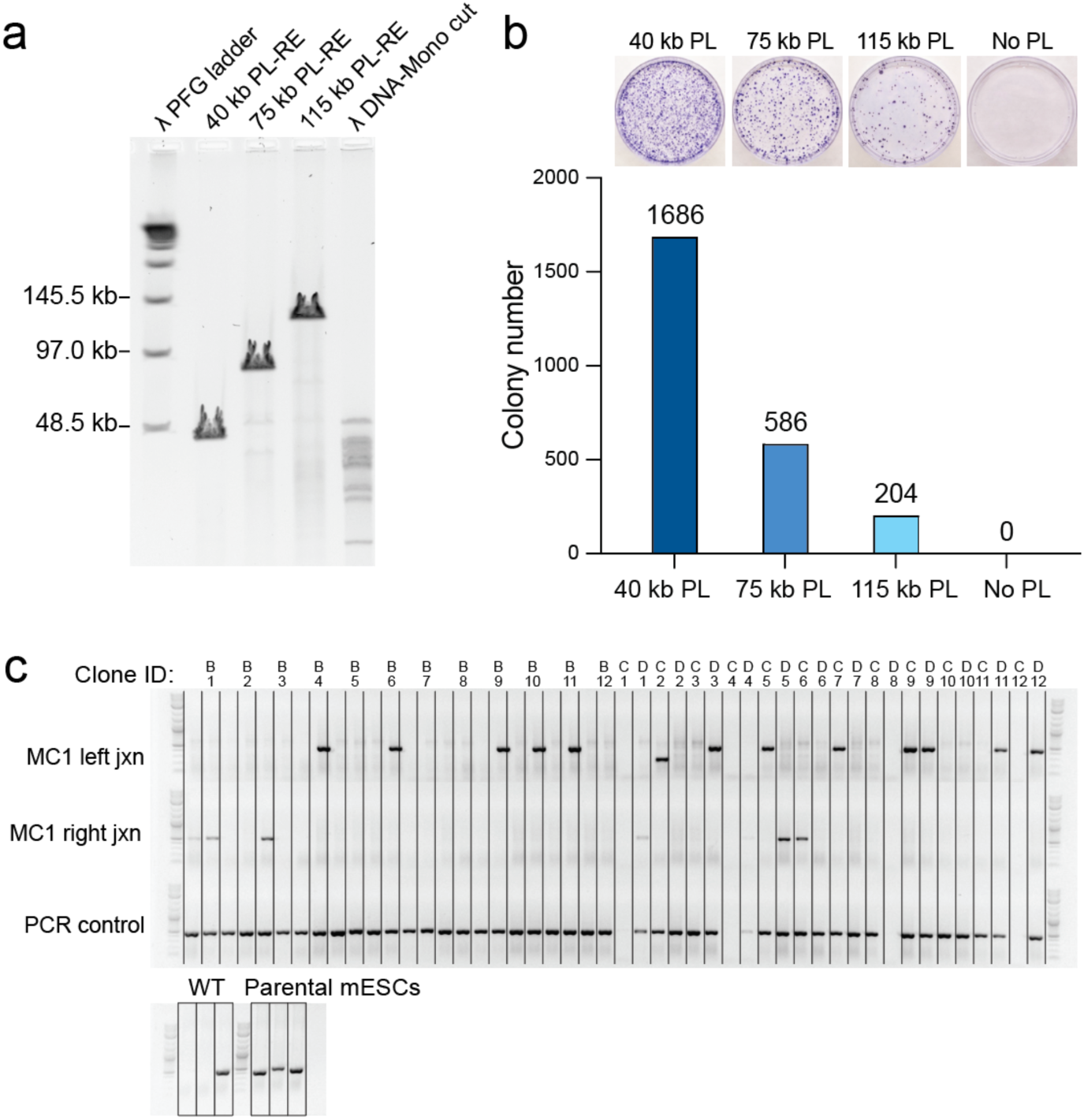
(*match to* Figure 2) Iterative genome writing with mSwAP-In. (**a**) Pulse field gel electrophoresis analysis of three *Trp53* downstream payloads linearized with a single-cutter. PL-RE, payload DNA digested with restriction enzyme. (**b**) Total colony number for the 40 kb, 75 kb and 115 kb payload deliveries using mSwAP-In. mESC colonies were fixed and stained with crystal violet. **(c)** Genotyping PCR of MC1 removal. Clone B1-C12 were from with-repair-donor group, clone D1-D12 were from without-repair-donor group. Jxn, junction PCR amplicon.

**Fig. S4.**
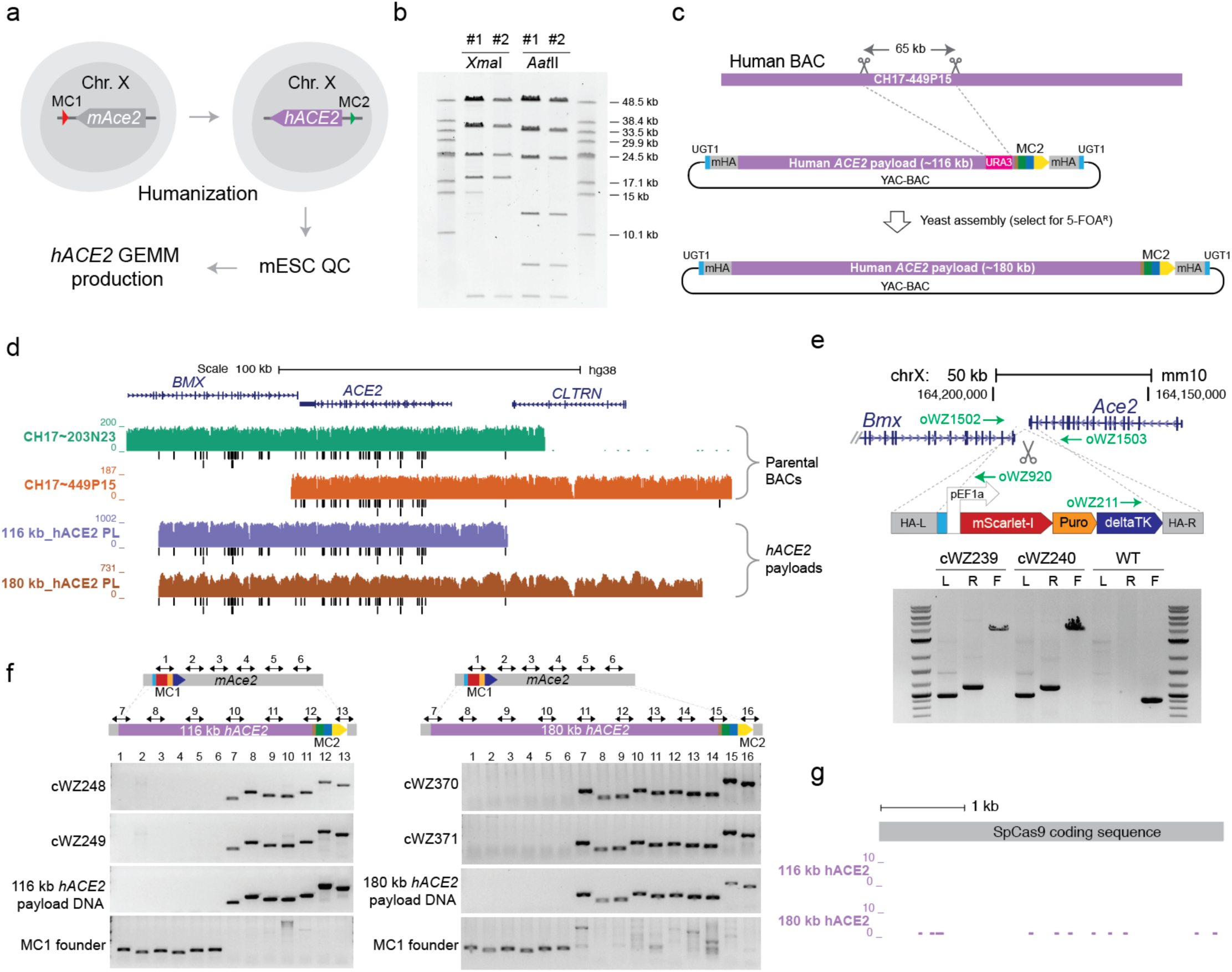
(*match to* Figure 3) Genomically rewriting mouse *Ace2* with human *ACE2*. (**a**) Schematic workflow. (**b**) Restriction enzyme digestion verification of 116 kb *hACE2* payload. Digestion products were separated using agarose gel by pulse field gel electrophoresis (see methods). (**c**) 180 kb *h*ACE2 payload assembly. Scissors mark *in vitro* CRISPR-Cas9 digestion sites. mHA, mouse homology arm. (**d**) Sequencing coverage of two human BACs and the two *hACE2* payloads mapped to hg38. Black bars represent SNPs. (**e**) MC1 integration downstream of *mAce2*. Integration was confirmed by junction PCR. HA-L, left homology arm, HA-R, right homology arm. L, left junction assay with primers oWZ1502 and oWZ920. R, right junction assay with primers oWZ211 and oWZ1503. F, full MC1 amplification with primers oWZ1502 and oWZ1503. (**f**) Genotyping PCR analysis of 116 kb and 180 kb *hACE2* mSwAP-In clones. Double headed arrows mark PCR amplicons from either m*Ace2* (assays 1-6) or *hACE2* payloads (assays 7-16). (**g**) Sequencing coverage of Cas9 in 116 kb and 180 kb *hACE2* mESCs.

**Fig. S5.**
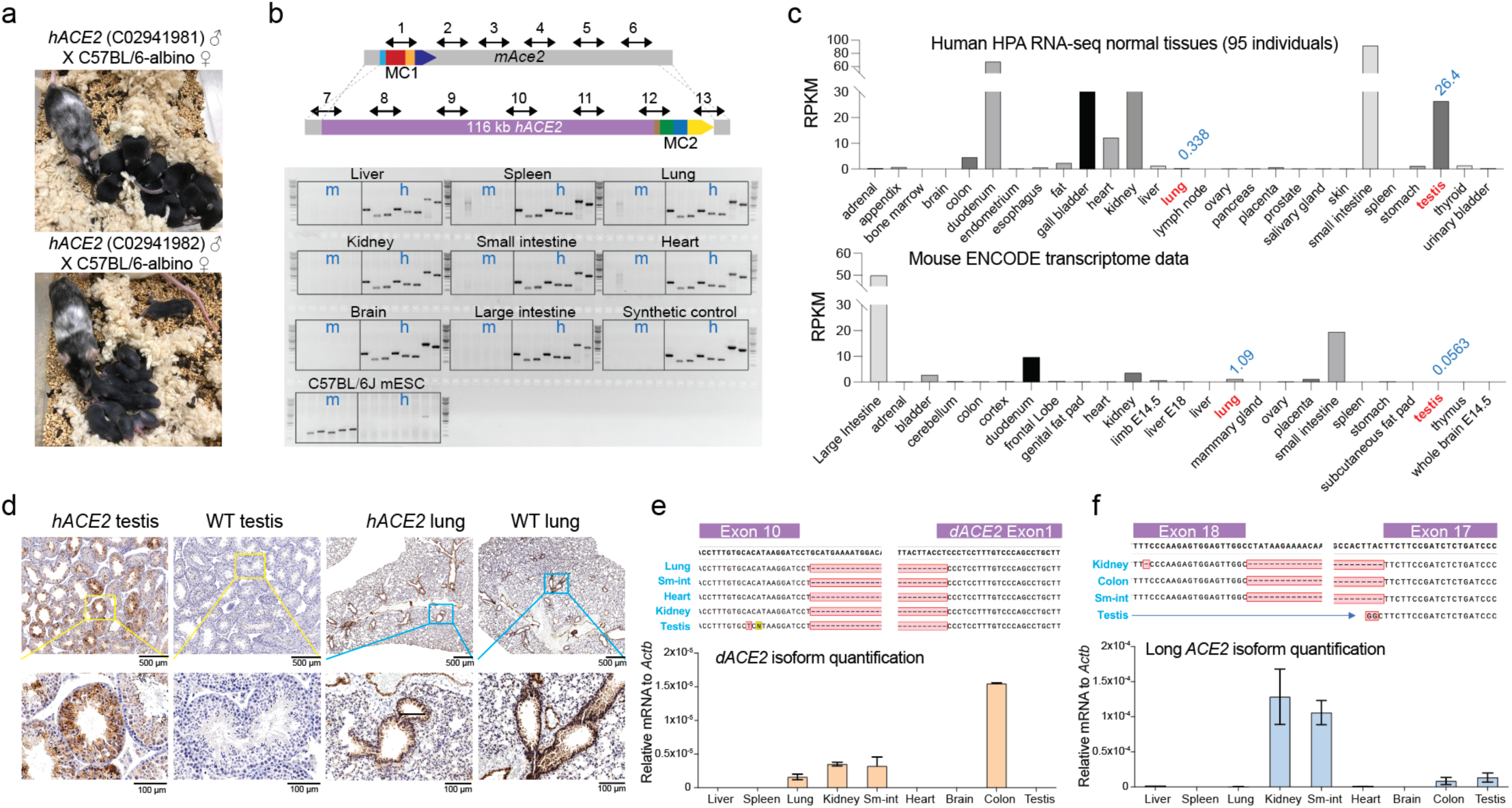
(*match to* Figure 4) *hACE2* expression characterization. (**a**) Two chimeric *hACE2* males showed 100% germline transmission rate when crossing with C57BL/6-albino females. (**b**) Genotyping PCR analysis of eight tissues from a tetraploid complementation-derived male. Double headed arrows are PCR amplicons from either mouse *Ace2* locus or human *ACE2* locus. m, *mAce2* amplicons; h, *hACE2* amplicons. (**c**) Human *ACE2* and mouse *Ace2* transcriptomic data from NCBI database. Lung and testis are highlighted in red with RPKM values indicated above. (**d**) Immunohistochemistry staining of testis and lung from *hACE2* and wild-type mice. An ACE2 antibody (Abcam, ab108209) that preferentially binds to human ACE2 was used for staining. Yellow and blue boxes are the magnified area. (**e-f**) Two *hACE2* isoforms detected in *hACE2* mice. *dACE2* novel junction Sanger sequencing analysis (up) and tissue distribution (down) (e). *hACE2* transcript 3 junction Sanger sequencing analysis (up) and tissue (down) (f). Expression levels were normalized to *Actb* gene, bars represent mean ± SD of three technical replicates.

**Fig. S6.**
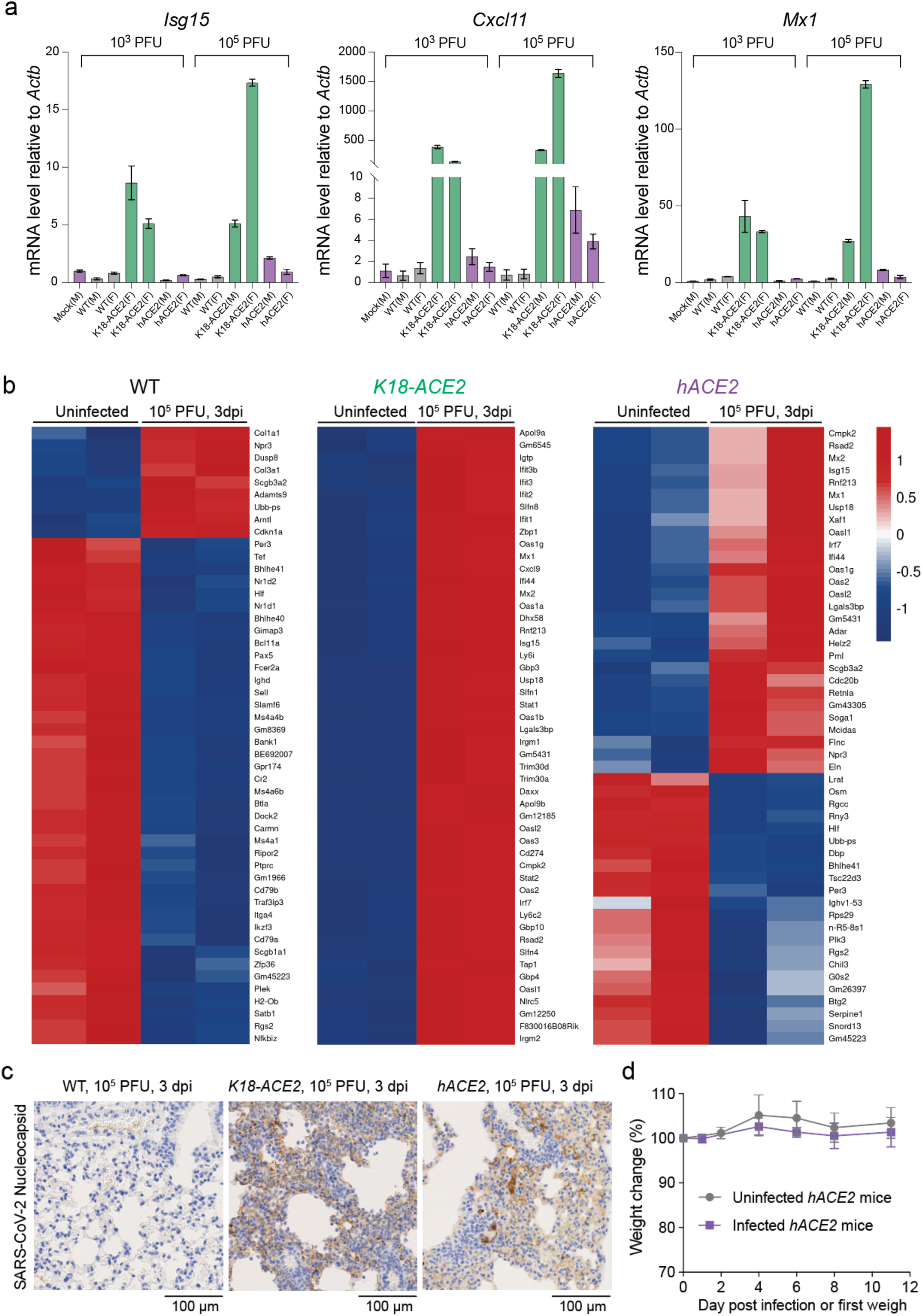
(*match to* Figure 5) SARS-CoV-2 infection of *hACE2* GEMM. **(a)** RT-qPCR analysis of three interferon-stimulated genes, *Isg15*, *Cxcl11*, *Mx1* in SARS-CoV-2 infected lungs at 3 dpi. Expression was normalized to *Actb* and to an uninfected control. Bars represent mean ± SD of three technical replicates. **(b)** Heatmaps of top 50 differentially expressed genes of wild-type, *K18-ACE2* and *hACE2* infected lungs comparing uninfected lungs. Color scale, z-score. **(c)** IHC staining of wild-type, *K18-ACE2* and *hACE2* lungs with SARS-CoV-2 nucleocapsid protein antibody (Thermo Fisher Scientific, MA1-7404). Female lung sections that are adjacent to H&E staining section (Fig. 5d) were used. **(d)** Weight curve comparison between SARS-CoV-2 infected (10^5^ PFU) and uninfected *hACE2* mice, n=4 in each group. Bars represent mean ± SD of biological replicates.

**Table S1.**
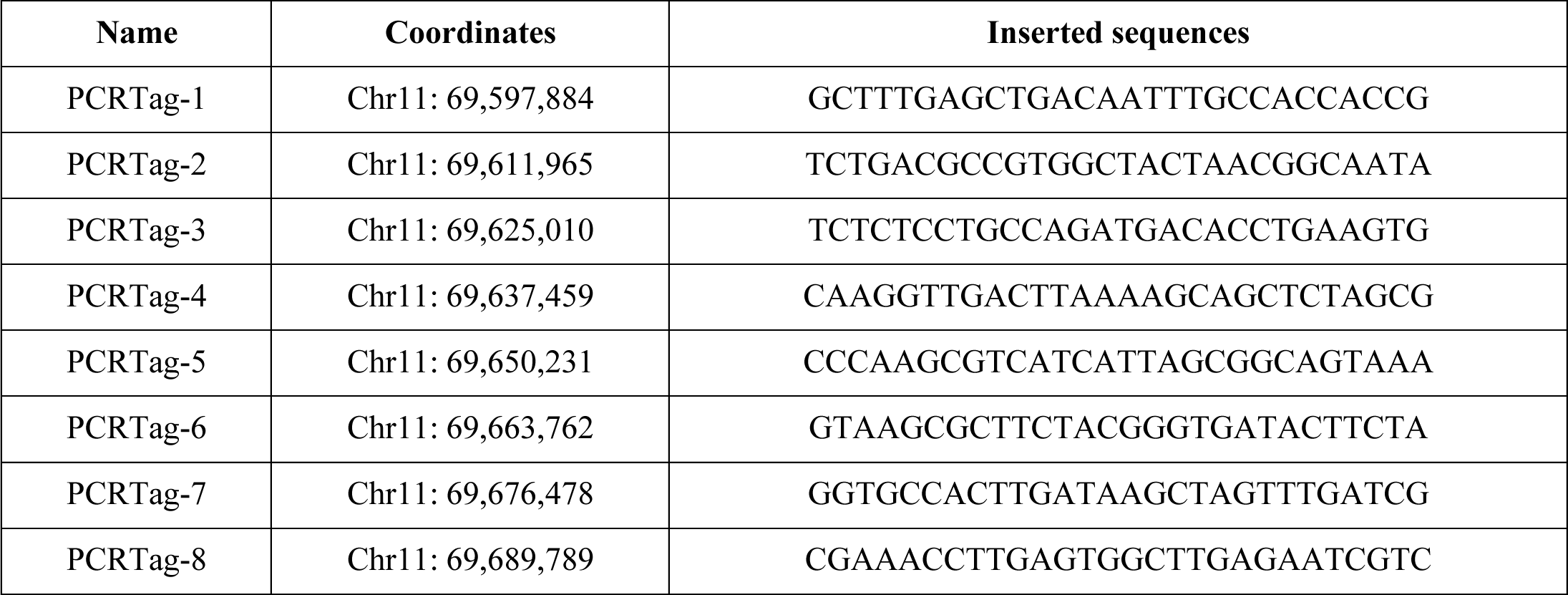
PCRTag sequences and insertion coordinates for the subsequent mSwAP-In of *synTrp53*.

**Table S2.** Summary of tetraploid blastocyst injection success rate.

| Tetraploid injected mESC clones | No. of Injected Embryos | No. of Pups | Birth Rate |
| --- | --- | --- | --- |
| 116 kb *hACE2*-cWZ271 | 25 | 5 | 20% |
| 116 kb *hACE2*-cWZ272 | 25 | 2 | 8% |
| 180 kb *hACE2*-cWZ329 | 25 | 5 | 20% |
| 180 kb *hACE2*-cWZ328 | 20 | 6 | 30% |
| 180 kb *hACE2*-cWZ323 | 25 | 5 | 20% |

**Table S3.**
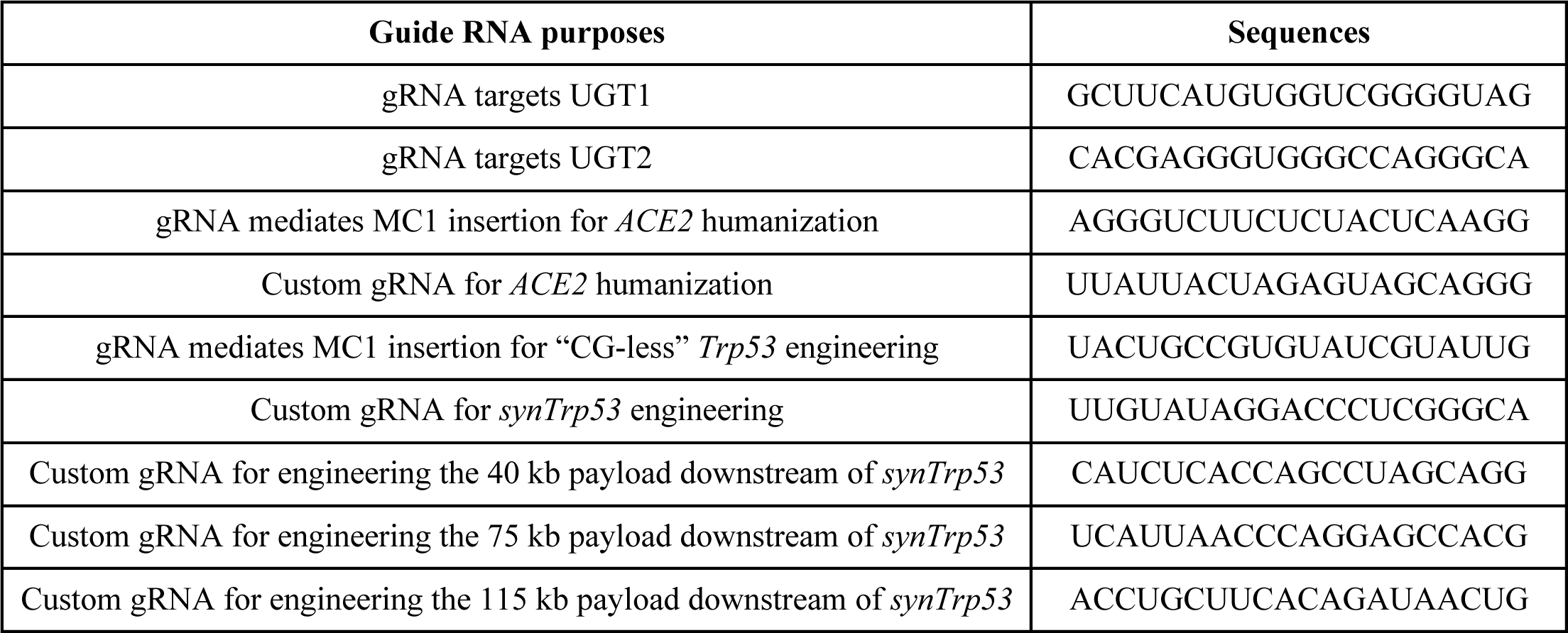
Guide RNA sequences.

**Table S4.**
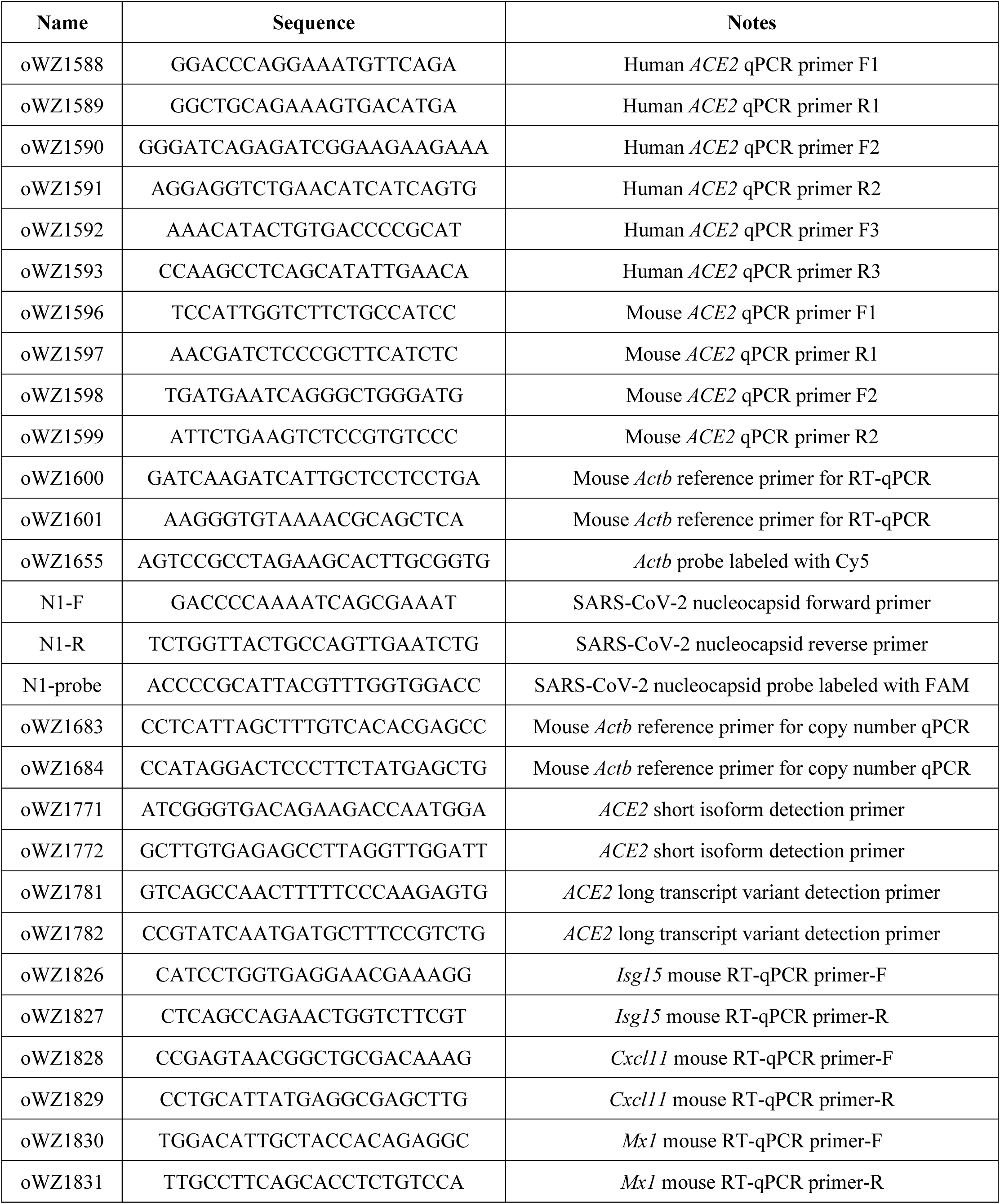
RT-qPCR primers and probes.

## References

1. Fredens, J. et al. Total synthesis of Escherichia coli with a recoded genome. Nature 569, 514–518 (2019).

2. Gibson, D. G., et al. Complete Chemical Synthesis, Assembly, and Cloning of a Mycoplasma genitalium Genome. Science 319, 1215–1220 (2008).

3. Gibson, D. G. et al. Creation of a bacterial cell controlled by a chemically synthesized genome. Science 329, 52–56 (2010).

4. Dymond, J. S. et al. Synthetic chromosome arms function in yeast and generate phenotypic diversity by design. Nature 477, 471–476 (2011).

5. Annaluru, N. et al. Total synthesis of a functional designer eukaryotic chromosome. Science 344, 55–58 (2014).

6. Richardson, S. M. et al. Design of a synthetic yeast genome. Science 355, 1040–1044 (2017).

7. Shen, Y. et al. Deep functional analysis of synII, a 770-kilobase synthetic yeast chromosome. Science 355, eaaf4791 (2017).

8. Xie, Z.-X. et al. “Perfect” designer chromosome V and behavior of a ring derivative. Science 355, eaaf4704 (2017).

9. Mitchell, L. A. et al. Synthesis, debugging, and effects of synthetic chromosome consolidation: synVI and beyond. Science 355, eaaf4831 (2017).

10. Wu, Y. et al. Bug mapping and fitness testing of chemically synthesized chromosome X. Science 355, eaaf4706 (2017).

11. Zhang, W. et al. Engineering the ribosomal DNA in a megabase synthetic chromosome. Science 355, (2017).

12. Dixon, J. R. et al. Topological domains in mammalian genomes identified by analysis of chromatin interactions. Nature 485, 376–380 (2012).

13. Wallace, H. A. C. et al. Manipulating the mouse genome to engineer precise functional syntenic replacements with human sequence. Cell 128, 197–209 (2007).

14. Brosh, R. et al. A versatile platform for locus-scale genome rewriting and verification. Proc. Natl. Acad. Sci. 118, (2021).

15. Iacovino, M. et al. Inducible Cassette Exchange: A Rapid and Efficient System Enabling Conditional Gene Expression in Embryonic Stem and Primary Cells. STEM CELLS 29, 1580–1588 (2011).

16. Mitchell, L. A. et al. De novo assembly and delivery to mouse cells of a 101 kb functional human gene. Genetics 218, (2021).

17. Nagy, A., Rossant, J., Nagy, R., Abramow-Newerly, W. & Roder, J. C. Derivation of completely cell culture-derived mice from early-passage embryonic stem cells. Proc. Natl. Acad. Sci. 90, 8424–8428 (1993).

18. Wang, Z.-Q., Kiefer, F., Urbánek, P. & Wagner, E. F. Generation of completely embryonic stem cell-derived mutant mice using tetraploid blastocyst injection. Mech. Dev. 62, 137–145 (1997).

19. Nagy, A. et al. Embryonic stem cells alone are able to support fetal development in the mouse. Development 110, 815– 821 (1990).

20. Eggan, K. et al. Hybrid vigor, fetal overgrowth, and viability of mice derived by nuclear cloning and tetraploid embryo complementation. Proc. Natl. Acad. Sci. U. S. A. 98, 6209–6214 (2001).

21. Zhang, Y. E. & Long, M. New genes contribute to genetic and phenotypic novelties in human evolution. Curr. Opin. Genet. Dev. 29, 90–96 (2014).

22. Anguita, E. et al. Deletion of the mouse alpha-globin regulatory element (HS-26) has an unexpectedly mild phenotype. Blood 100, 3450–3456 (2002).

23. Engle, S. J. et al. HPRT-APRT-deficient mice are not a model for lesch-nyhan syndrome. Hum. Mol. Genet. 5, 1607– 1610 (1996).

24. McCray, P. B. et al. Lethal Infection of K18-hACE2 Mice Infected with Severe Acute Respiratory Syndrome Coronavirus. J. Virol. 81, 813–821 (2007).

25. Moore, J. E. et al. Expanded encyclopaedias of DNA elements in the human and mouse genomes. Nature 583, 699–710 (2020).

26. Tam, V. et al. Benefits and limitations of genome-wide association studies. Nat. Rev. Genet. 20, 467–484 (2019).

27. Antoch, M. P. et al. Functional identification of the mouse circadian Clock gene by transgenic BAC rescue. Cell 89, 655– 667 (1997).

28. Mendez, M. J. et al. Functional transplant of megabase human immunoglobulin loci recapitulates human antibody response in mice. Nat. Genet. 15, 146–156 (1997).

29. Lamb, B. T. et al. Introduction and expression of the 400 kilobase amyloid precursor protein gene in transgenic mice [corrected]. Nat. Genet. 5, 22–30 (1993).

30. Macdonald, L. E. et al. Precise and in situ genetic humanization of 6 Mb of mouse immunoglobulin genes. Proc. Natl. Acad. Sci. 111, 5147–5152 (2014).

31. Murphy, A. J. et al. Mice with megabase humanization of their immunoglobulin genes generate antibodies as efficiently as normal mice. Proc. Natl. Acad. Sci. U. S. A. 111, 5153–5158 (2014).

32. Dinnon, K. H. et al. A mouse-adapted model of SARS-CoV-2 to test COVID-19 countermeasures. Nature 586, 560–566 (2020).

33. Leist, S. R. et al. A Mouse-Adapted SARS-CoV-2 Induces Acute Lung Injury and Mortality in Standard Laboratory Mice. Cell 183, 1070–1085.e12 (2020).

34. Shuai, H. et al. Emerging SARS-CoV-2 variants expand species tropism to murines. eBioMedicine 73, (2021).

35. Imai, M., et al. Characterization of a new SARS-CoV-2 variant that emerged in Brazil. Proc. Natl. Acad. Sci. 118, e2106535118 (2021).

36. Tseng, C.-T. K. et al. Severe Acute Respiratory Syndrome Coronavirus Infection of Mice Transgenic for the Human Angiotensin-Converting Enzyme 2 Virus Receptor. J. Virol. 81, 1162–1173 (2007).

37. Yang, X.-H. et al. Mice transgenic for human angiotensin-converting enzyme 2 provide a model for SARS coronavirus infection. Comp. Med. 57, 450–459 (2007).

38. Menachery, V. D. et al. SARS-like WIV1-CoV poised for human emergence. Proc. Natl. Acad. Sci. 113, 3048–3053 (2016).

39. Onabajo, O. O. et al. Interferons and viruses induce a novel truncated ACE2 isoform and not the full-length SARS-CoV-2 receptor. Nat. Genet. 52, 1283–1293 (2020).

40. Boeke, J. D., et al. GENOME ENGINEERING. The Genome Project-Write. Science 353, 126–127 (2016).

41. van der Lugt, N., Maandag, E. R., te Riele, H., Laird, P. W. & Berns, A. A pgk::hprt fusion as a selectable marker for targeting of genes in mouse embryonic stem cells: disruption of the T-cell receptor delta-chain-encoding gene. Gene 105, 263–267 (1991).

42. Chandran, S. & Shapland, E. Efficient Assembly of DNA Using Yeast Homologous Recombination (YHR). in Synthetic DNA: Methods and Protocols (ed. Hughes, R. A.) 187–192 (Springer, 2017). doi:10.1007/978-1-4939-6343-0_14.

43. Li, X., et al. piggyBac transposase tools for genome engineering. Proc. Natl. Acad. Sci. 110, E2279–E2287 (2013).

44. Brosh, R. & Rotter, V. When mutants gain new powers: news from the mutant p53 field. Nat. Rev. Cancer 9, 701–713 (2009).

45. Baugh, E. H., Ke, H., Levine, A. J., Bonneau, R. A. & Chan, C. S. Why are there hotspot mutations in the TP53 gene in human cancers? Cell Death Differ. 25, 154–160 (2018).

46. Freed-Pastor, W. A. & Prives, C. Mutant p53: one name, many proteins. Genes Dev. 26, 1268–1286 (2012).

47. Shen, J. C., Rideout, W. M. & Jones, P. A. The rate of hydrolytic deamination of 5-methylcytosine in double-stranded DNA. Nucleic Acids Res. 22, 972–976 (1994).

48. Barnes, D. E. & Lindahl, T. Repair and genetic consequences of endogenous DNA base damage in mammalian cells. Annu. Rev. Genet. 38, 445–476 (2004).

49. Chen, J. X., Zheng, Y., West, M. & Tang, M. S. Carcinogens preferentially bind at methylated CpG in the p53 mutational hot spots. Cancer Res. 58, 2070–2075 (1998).

50. Yoon, J. H. et al. Methylated CpG dinucleotides are the preferential targets for G-to-T transversion mutations induced by benzo[a]pyrene diol epoxide in mammalian cells: similarities with the p53 mutation spectrum in smoking-associated lung cancers. Cancer Res. 61, 7110–7117 (2001).

51. Denissenko, M. F., Pao, A., Tang, M. & Pfeifer, G. P. Preferential formation of benzo[a]pyrene adducts at lung cancer mutational hotspots in P53. Science 274, 430–432 (1996).

52. He, H. et al. p53 and p73 Regulate Apoptosis but Not Cell-Cycle Progression in Mouse Embryonic Stem Cells upon DNA Damage and Differentiation. Stem Cell Rep. 7, 1087–1098 (2016).

53. Rausch, T., et al. DELLY: structural variant discovery by integrated paired-end and split-read analysis. Bioinforma. Oxf. Engl. 28, i333–i339 (2012).

54. Alexander, M. R. et al. Predicting susceptibility to SARS-CoV-2 infection based on structural differences in ACE2 across species. FASEB J. 34, 15946–15960 (2020).

55. Yan, R. et al. Structural basis for the recognition of SARS-CoV-2 by full-length human ACE2. Science 367, 1444–1448 (2020).

56. Li, W. et al. Efficient Replication of Severe Acute Respiratory Syndrome Coronavirus in Mouse Cells Is Limited by Murine Angiotensin-Converting Enzyme 2. J. Virol. 78, 11429–11433 (2004).

57. Kumari, P. et al. Neuroinvasion and Encephalitis Following Intranasal Inoculation of SARS-CoV-2 in K18-hACE2 Mice. Viruses 13, 132 (2021).

58. Badawi, S. & Ali, B. R. ACE2 Nascence, trafficking, and SARS-CoV-2 pathogenesis: the saga continues. Hum. Genomics 15, 8 (2021).

59. Valenzuela, D. M. et al. High-throughput engineering of the mouse genome coupled with high-resolution expression analysis. Nat. Biotechnol. 21, 652–659 (2003).

60. Sun, S.-H. et al. A Mouse Model of SARS-CoV-2 Infection and Pathogenesis. Cell Host Microbe 28, 124–133.e4 (2020).

61. Sefik, E. et al. A humanized mouse model of chronic COVID-19. Nat. Biotechnol. 1–15 (2021) doi:10.1038/s41587-021-01155-4.

62. Ma, X. et al. Pathological and molecular examinations of postmortem testis biopsies reveal SARS-CoV-2 infection in the testis and spermatogenesis damage in COVID-19 patients. Cell. Mol. Immunol. 18, 487–489 (2021).

63. Fan, C., Lu, W., Li, K., Ding, Y. & Wang, J. ACE2 Expression in Kidney and Testis May Cause Kidney and Testis Infection in COVID-19 Patients. Front. Med. 7, (2021).

64. Ziegler, C. G. K. et al. SARS-CoV-2 Receptor ACE2 Is an Interferon-Stimulated Gene in Human Airway Epithelial Cells and Is Detected in Specific Cell Subsets across Tissues. Cell 181, 1016–1035.e19 (2020).

65. Hoagland, D. A. et al. Leveraging the antiviral type I interferon system as a first line of defense against SARS-CoV-2 pathogenicity. Immunity 54, 557–570.e5 (2021).

66. Sia, S. F. et al. Pathogenesis and transmission of SARS-CoV-2 in golden hamsters. Nature 583, 834–838 (2020).

67. Roberts, A. et al. Severe acute respiratory syndrome coronavirus infection of golden Syrian hamsters. J. Virol. 79, 503– 511 (2005).

68. Iwatsuki-Horimoto, K. et al. Syrian Hamster as an Animal Model for the Study of Human Influenza Virus Infection. J. Virol. 92, e01693–17 (2018).

69. Suresh, V. et al. Tissue Distribution of ACE2 Protein in Syrian Golden Hamster (Mesocricetus auratus) and Its Possible Implications in SARS-CoV-2 Related Studies. Front. Pharmacol. 11, (2021).

70. Lee, I. T. et al. ACE2 localizes to the respiratory cilia and is not increased by ACE inhibitors or ARBs. Nat. Commun. 11, 5453 (2020).

71. Schaefer, I.-M. et al. In situ detection of SARS-CoV-2 in lungs and airways of patients with COVID-19. Mod. Pathol. 33, 2104–2114 (2020).

72. Rhie, A. et al. Towards complete and error-free genome assemblies of all vertebrate species. Nature 592, 737–746 (2021).

73. Nurk, S. et al. The complete sequence of a human genome. Science 376, 44–53 (2022).

